# Performance of Core Genome Multilocus Sequence Typing Compared to Capillary-Electrophoresis PCR Ribotyping and SNP Analysis of *Clostridioides difficile*

**DOI:** 10.1101/2021.08.10.455895

**Authors:** A Baktash, J Corver, C Harmanus, WK Smits, W Fawley, MH Wilcox, N Kumar, DW Eyre, A Indra, A Mellmann, EJ Kuijper

## Abstract

*Clostridioides difficile* is the most common cause of antibiotic-associated gastrointestinal infections. Capillary-electrophoresis (CE)-PCR ribotyping is currently the gold standard for *C. difficile* typing but lacks discriminatory power to study transmission and outbreaks in detail. New molecular methods have the capacity to differentiate better, but backward compatibility with CE-PCR ribotyping must be assessed. Using a well-characterized collection of diverse strains (N=630; 100 unique ribotypes [RTs]), we aimed to investigate PCR ribotyping prediction from core genome multilocus sequence typing (cgMLST). Additionally, we compared the discriminatory power of cgMLST (SeqSphere & EnteroBase) and whole genome MLST (wgMLST) (EnteroBase) with single nucleotide polymorphism (SNP) analysis). A unique cgMLST profile (>6 allele differences) was observed in 82/100 ribotypes, indicating sufficient backward compatibility. Intra-RT allele difference varied per ribotype and MLST clade. Application of cg/wgMLST and SNP analysis in two outbreak settings with ribotypes RT078 and RT181 (known with a low intra-ribotype allele difference) showed no distinction between outbreak- and non-outbreak strains, in contrast to wgMLST and SNP analysis. We conclude that cgMLST has the potential to be an alternative to CE-PCR ribotyping. The method is reproducible, easy to standardize and offers higher discrimination. However, in some ribotype complexes adjusted cut-off thresholds and epidemiological data are necessary to recognize outbreaks. We propose to decrease the current threshold of 6 to 3 alleles to better identify outbreaks.

## INTRODUCTION

*Clostridioides difficile* is a Gram-positive anaerobic bacterium that is associated with nosocomial gastrointestinal infection (1) (2). It is estimated that there were almost 500,000 patients with *C. difficile* infection (CDI) and around 29,000 deaths in the United States in 2011 (2). Individuals with *C. difficile* infection (CDI) are an important source of *C. difficile* transmission in healthcare settings (2). Typing of *C. difficile* is necessary for infection control, epidemiology and evaluation of treatment. Several methods are used for typing *C. difficile*, including capillary electrophoresis (CE) PCR ribotyping (3) (4) and multilocus sequence typing (MLST) (5). CE-PCR ribotyping is currently the gold standard. However, it does not provide sufficient discriminatory power to distinguish related strains (6). Furthermore, for CE-PCR ribotyping, standardization and interlaboratory comparisons are difficult to establish (7), whereas for MLST this is relatively simple. In the case of a suspected outbreak CE-PCR ribotyping can be used in combination with multilocus variable-number tandem repeat (VNTR) analysis (MLVA) for subtyping of strains belonging to one PCR ribotype (8). This combination of methods is usually sufficient to type strains and understand transmission events. However, these methods do not provide sufficient information about strain characteristics (e.g. possession of virulence and resistance genes) and possible treatment failures (relapse vs. reinfection). The techniques are also less suitable to study transmission and to determine the role of symptomatic and asymptomatic patients in hospital acquired CDI (9). Therefore, typing methods with more discriminatory power and preferably based on better standardized whole genome sequencing (WGS) are urgently needed.

There are two commonly applied methods to identify genomic variations using WGS. Single nucleotide polymorphism (SNP) analysis usually uses a reference genome and detects SNPs between the reference genome and the studied genome (10). SNP analysis provides the highest resolution, but it is relatively slow, requires extensive bioinformatic tools, is difficult to standardize and typing nomenclature is missing (11), (12), (9). The second approach is based on gene-by-gene allelic profiling of the core genome (cgMLST) or whole genome (wgMLST) (13). cgMLST provides high discriminatory power, is more rapid than SNP analysis, offers reasonably accurate reproducibility (11) and could be used as a typing method since the scheme is maintained by a centralized database (14).

Currently there are several cg/wgMLST schemes available for *C. difficile*, both commercially and publicly. The first commercial platform is SeqSphere+ software [Ridom GmbH, Germany] comprising of a scheme (the cgMLST.org Nomenclature Server) using up to 2147 core genes and 1357 accessory genes out of 3756 genes present in strain 630 (14). The second is BioNumerics [bioMérieux, France] with the cgMLST/wgMLST scheme developed by Applied-Maths, comprising 1999 core genes and 6713 accessory genes and several other genes associated with virulence, antimicrobial resistance and others from different *C. difficile* strains (15). Besides these 2 commercial platforms, there is a publicly available cg/wgMLST scheme from EnteroBase [University of Warwick, UK] consisting of 2556 genes for the cgMLST scheme and up to 13763 genes for the wgMLST scheme (16). The cgMLST scheme of EnteroBase (EB cgMLST) is also available through the Center for Genomic Epidemiology (cgMLSTFinder 1.1; https://cge.cbs.dtu.dk/services/cgMLSTFinder/).

Several studies have been published on the application of cgMLST (14), (11), (15), (16). Most studies show that cgMLST is backward compatible with CE-PCR ribotyping but only a restricted number of different ribotypes were analysed and outbreaks were not included. Recently, Seth-Smith and colleagues showed that cgMLST predicted 36 ribotypes using nearly 300 well characterised clinical strains from Switzerland. However, some ribotypes complexes (RT 078/126) has a low genomic difference, whereas other ribotypes (e.g., RT 023) were very disperse (17). Our study builds upon previous work by assessing backward compatibility more in depth, using 100 unique ribotypes and changing thresholds to determine optimal differentiation between ribotypes. Furthermore, we analyse the performance of CE-PCR ribotyping, cgMLST, wgMLST and SNP analysis by using multiple software programs (SeqSphere & EnteroBase) and applied the methods on two outbreaks. Importantly, our study shows that a threshold of ≤ 3 targets/alleles is needed for *C. difficile* isolates that are likely to belong to the same clone in an outbreak setting.

## MATERIALS AND METHODS

### Sequence data

The NCBI database was searched at the start of this study for sequenced closed *C. difficile* genomes, this resulted in 4845 available genomes. Only sequence data generated on Illumina sequencing platform and representing known ribotypes were selected. A random selection of overrepresented strains (e.g. RT027 and RT078) were included. This approach resulted in 609 complete genome sequences that were analysed. Besides downloaded strains from the NCBI database we included also 21 recently sequenced strains at Leiden University Medical Center (LUMC). This comprised fifteen Greek RT181 CDI outbreak strains that were already sequenced for a previous study (PRJEB36956, Table S1, (18) and 6 strains from a Dutch CDI outbreak due to RT078. For sequencing of strains, total DNA was isolated from cultured bacteria. A few colonies were emulsified in Tris/EDTA (TE) buffer and heated at 100° C for 10 minutes according to the Griffiths *et al*. protocol (5). DNA was sequenced at Genome Scan B.V., Leiden, The Netherlands, on an Illumina NovaSeq 6000 after preparation with the NebNext Ultra II DNA library prep kit for Illumina. This produced on average 3 million paired-end reads (read size 150bp) per sample, with a minimum of 90% reads with a quality of ≥30.

### Ridom cgMLST

Ridom^®^ SeqSphere^+^ (version 6.0.2; Ridom GmbH, Münster, Germany) was run with default settings for quality trimming, *de novo* assembly and allele calling on a Microsoft Windows operating system. Quality trimming occurred at both 5’-ends and 3’-end until an average base quality of 30 was reached (length of 20 bases and a 120-fold coverage) (14), (13). *De novo* assembly was performed using the SKESA assembler version 2.3.0 (19) integrated in SeqSphere^+^ (20) using default settings for SKESA. SeqSphere^+^ scanned for the defined genes using BLAST (21) with criteria described previously (22), (13). For further analysis, distance matrices, minimum spanning trees and neighbour joining trees were constructed using the integrated features within SeqSphere^+^ with “pairwise ignoring missing values” option turned on.

### EnteroBase cgMLST and wgMLST

cgMLST was performed using cgMLST Finder 1.1, available through the Center for Genomic Epidemiology (cgMLSTFinder 1.1; https://cge.cbs.dtu.dk/services/cgMLSTFinder/). Genomic data was processed using automated pipelines inside EnteroBase, as described in detail previously (23). In short, *de novo* assembly of Illumina sequence reads was performed using Spades v3.10 (24). In order to pass quality control, assemblies were needed to comply with criteria described previously (16). BLASTn and UBLASTP were used to align assemblies to alleles. EnteroBase module MLSType was used to assess allele numbers and cluster types (23). cgMLST Finder 1.1 provides a distance matrix for analysis. Distance matrices were used to calculate the mean intra- and inter-allelic distance between different CE-PCR ribotypes. For wgMLST analysis, an *ad hoc* scheme was used based on the wgMLST scheme from EnteroBase (EB wgMLST) (16), (25). This *ad hoc* scheme was integrated in Ridom^®^ SeqSphere (14). *De novo* assembly, allele calling and further analysis were carried out as mentioned previously (under Ridom cgMLST).

### SNP analysis

SNPs were identified as previously described (26) using the webtool at the following address: http://cge.cbs.dtu.dk/services/CSIPhylogeny/. Default settings were used for the SNP analysis. *C. difficile* strain 630 (NC_009089) was used as the reference genome for all analyses. In short, reads were mapped to the reference sequence using BWA (version 0.7.2) (27). Depth at each position was calculated using genomeCoverageBed, which is a component of BEDTools (version 2.16.2) (28). SNPs were called using mpileup, which is a component of SAMTools (version 0.1.18) (29). Mapping quality (minimum of 25 reads) and SNP quality (SNPs were filtered out if quality was below 30 or if they were called within the vicinity of 10 bp of another SNP) were calculated by BWA and SAMTools, respectively. CSIPhylogeny 1.4 provides a distance matrix for analysis. Distance matrices (based on pairwise comparison, missing data were excluded) were used to calculate the mean intra- and inter-RT SNP distance between different CE-PCR ribotypes.

### Mean intra-ribotype allele difference

Mean intra-ribotype allele difference was determined for 19 ribotypes using distance matrices produced with cgMLST and wgMLST schemes and SNP analysis. From each ribotype, 3 to 13 strains were included. To prevent inclusion of related strains, e.g. from outbreak reports, we selected ribotypes with at least 3 strains from different geographic locations and/or from different collection years.

### Data availability

All own genome sequence data generated as part of this study were submitted to the NCBI/ENA under study number PRJEB46469. Sequence Read Archive (SRA) accession numbers for other analyzed genomes are provided in Table S1.

## RESULTS

### Ridom cgMLST is backward compatible with CE-PCR ribotyping

To test the backward compatibility of cgMLST (SeqSphere) with CE-PCR ribotyping, we compared cgMLST and CE-PCR ribotyping using a selection of sequenced *C. difficile* strains with known ribotypes (Janezic & Rupnik, 2019). Figure 1 depicts a neighbour joining tree based on the Ridom SeqSphere cgMLST scheme (SqSp cgMLST) including 100 different PCR ribotypes from all 5 MLST Clades. Most ribotypes show a different allelic profile in cgMLST in comparison with other ribotypes. However, there are ribotypes within every MLST clade that show low allele difference (<6 alleles) in comparison with other ribotypes.

**Figure 1:**
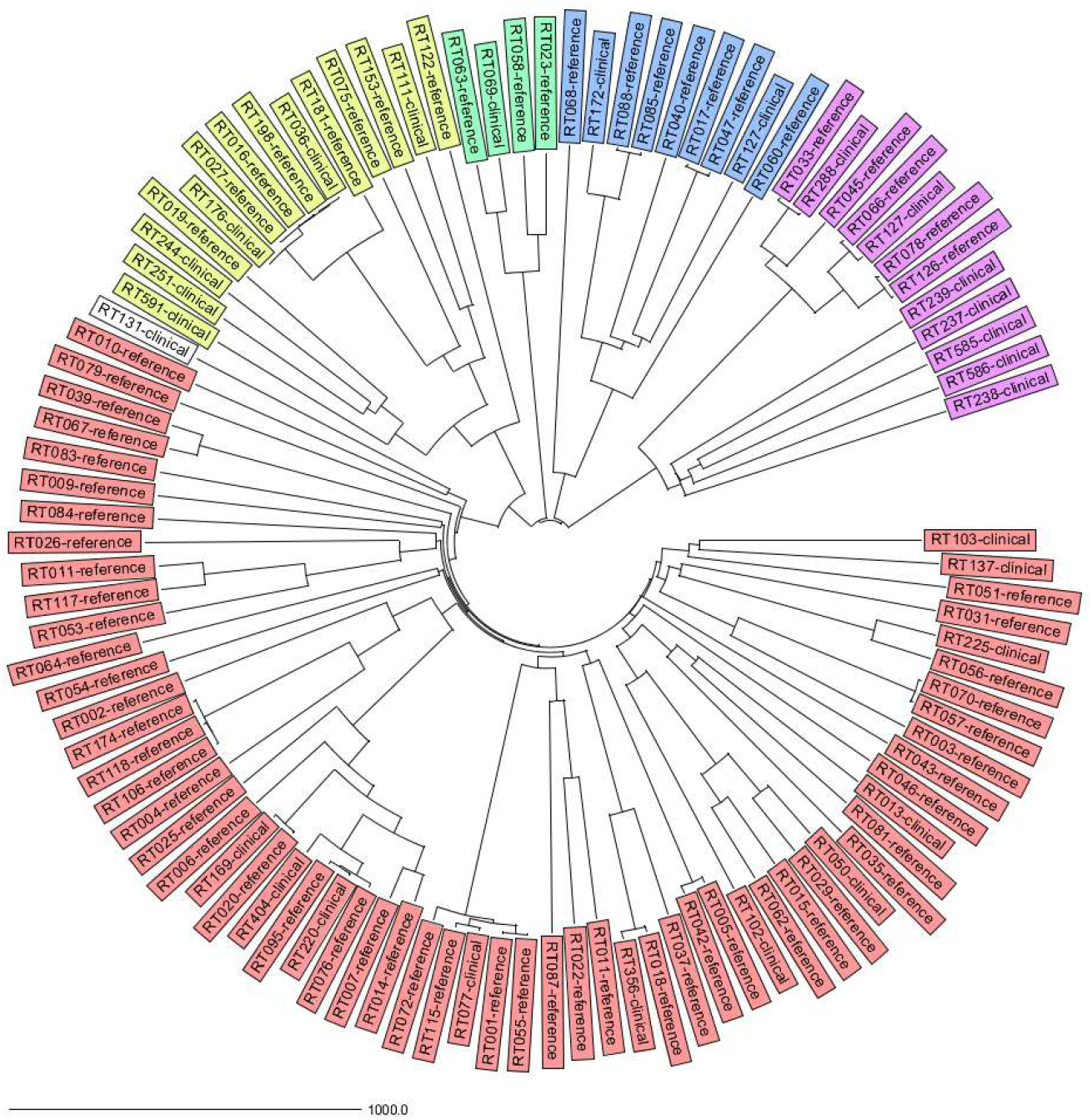
Neighbor joining tree from 100 unique ribotypes based on SqSp cgMLST allele difference. Each ribotype is depicted with “RTn” followed by “reference” (belonging to the Leeds-Leiden collection) or clinical (non-Leeds-Leiden strain). Ribotypes from MLST Clade 1, 2, 3, 4, 5 are colored red, yellow, green, blue and purple, respectively. RT131 stated as CD131-01, 131, has no designated MLST Clade and is shown in white. The distance is given in absolute allelic difference.

When all included strains (n=630 strains) from 100 unique ribotypes were analysed (shown in Table 1), 82 ribotypes were distinguishable, i.e., the strains within these ribotypes differed by >6 alleles from strains within other ribotypes. Eighteen ribotypes (18%) clustered together with 1-3 other ribotypes from the same clade and had ≤ 6 allelic differences. This was observed in Clades 1, 2 and 5. In figure 2 we show the ribotypes in each cluster and how these clusters vary at different thresholds (0-6 allelic differences). When the threshold was lowered from 6 to 0, the number of different ribotypes that clustered decreased from 13 to 2 (RT045 and RT127). The amount of clusters decreased from 14 to 1. Even at a threshold of 0 allele difference, these ribotypes showed clustering, demonstrating the limitation of short-read sequencing and cgMLST.

**Figure 2:**
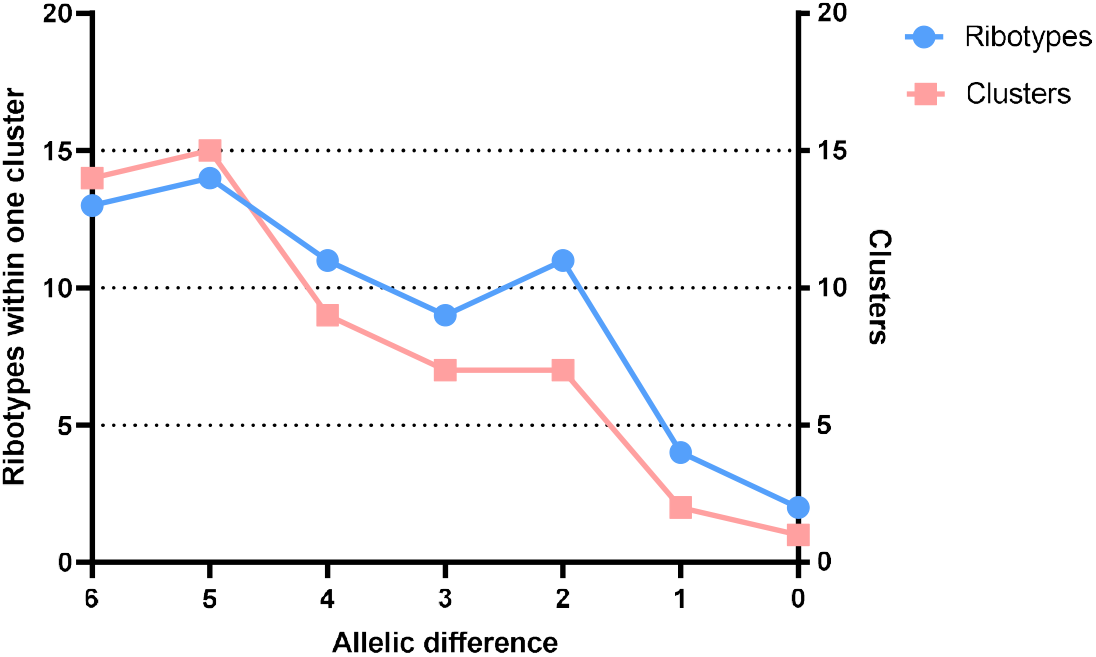
Clustering of different PCR ribotypes at different thresholds (0-6 allelic difference). The number of clustering PCR ribotypes is shown in blue and the amount of clusters at every threshold is shown in pink.

**Table 1:**
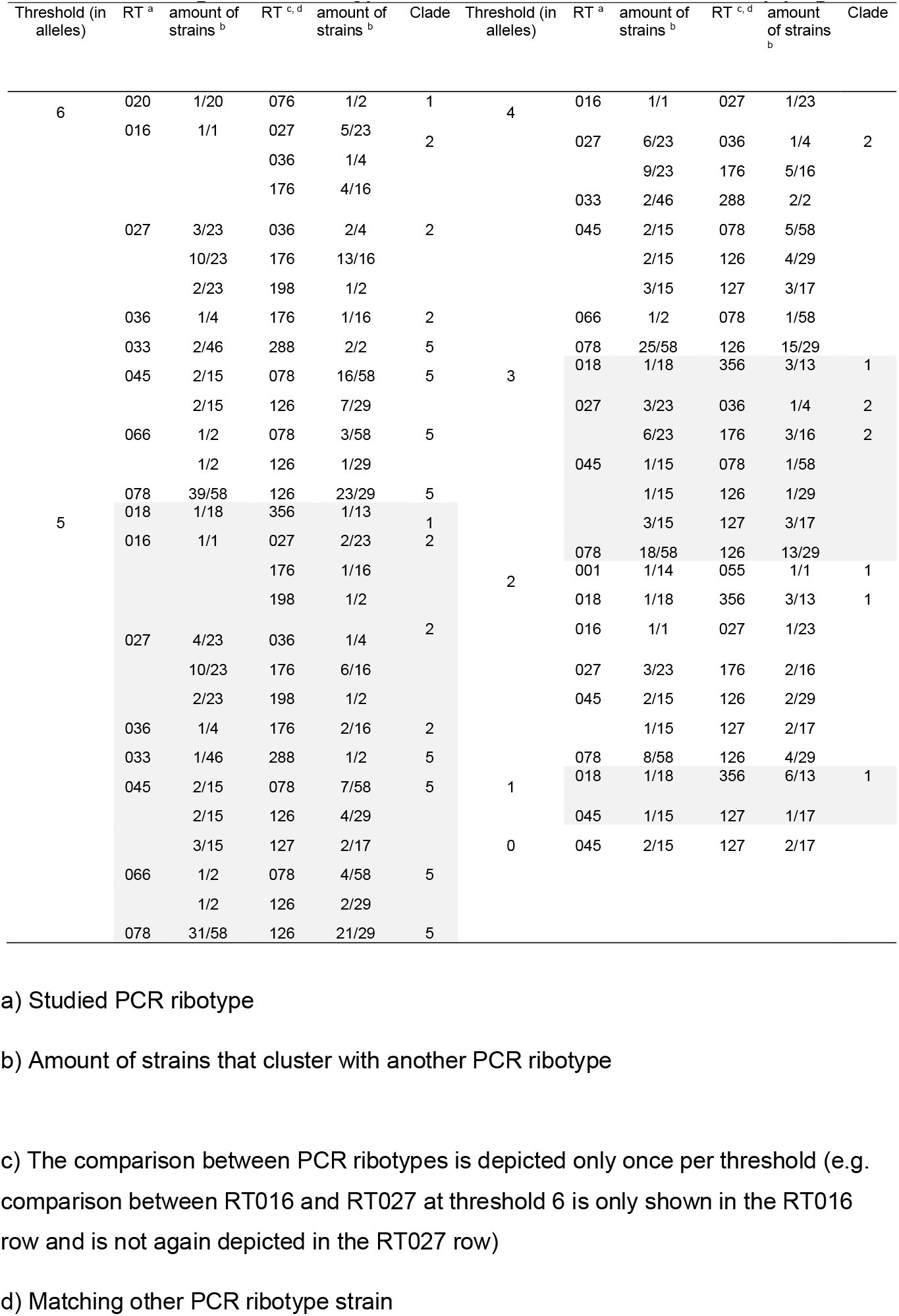
Clustering between ribotypes at different thresholds based on SqSp cgMLST.

### Intra-ribotype allele difference varies per ribotype and per MLST clade

We determined the mean allelic difference between strains from the same ribotype and tested if intra-ribotype allele differences vary between MLST clades and ribotypes. We also compared the mean intra-ribotype allele or SNP differences with cgMLST, wgMLST and SNP analysis. Mean intra-ribotype allele difference varied between ribotypes (Figure 3A). The method with the smallest scheme (SqSp cgMLST) showed the lowest intra-ribotype allele difference average (mean range of 5-376 alleles) whereas SNP analysis showed the highest average (mean range of 67-2563 alleles). As a comparison, the mean and median of the inter-ribotype allele difference were 1742 and 2131 alleles with SqSp cgMLST, respectively. Figure 3 A also shows that RT027 had an intra ribotype allele difference of 8.4 (SqSp cgMLST), 10.7 (EB cgMLST), 18.1 (EB wgMLST) and 100.7 (SNP). Another complex ribotype, RT078 showed 13.2, 15.5, 29.3 and 139.4, respectively. The most frequently found ribotype in Europe, RT014 showed 148.1, 173, 258.8 and 855.7 respectively. RT023 (clade 3) showed 108.7, 121.3, 157.5 and 1014.7, respectively. RT017 (clade 4) showed 22.3, 23.5, 63.7 and 129.3, respectively. EB wgMLST and SNP analysis showed similar results as cgMLST, but showed much higher average intra-ribotype allele and SNP difference. The ribotype with lowest intra-ribotype allele difference for clade 1 was again RT002 (64 alleles and 140 SNPs) and the highest was RT056 (650 alleles and 2563 SNPs). The ribotype with the lowest intra-ribotype difference from clade 2 was RT181 (11 alleles and 67 SNPs), whereas the highest was RT036 (39 alleles and 120 SNPs). RT023 from clade 3 showed an average of 158 intra-ribotype allele difference and 1015 SNP difference. RT017 from clade 4 showed 64 allele and 129 SNP difference. Lastly, RT126 from clade 5 showed the lowest difference (18 allele and 130 SNP differences) and RT127 the highest (379 allele and 592 SNP differences). SNP analysis showed the highest resolution and often >2 times difference in comparison with wgMLST.

**Figure 3:**
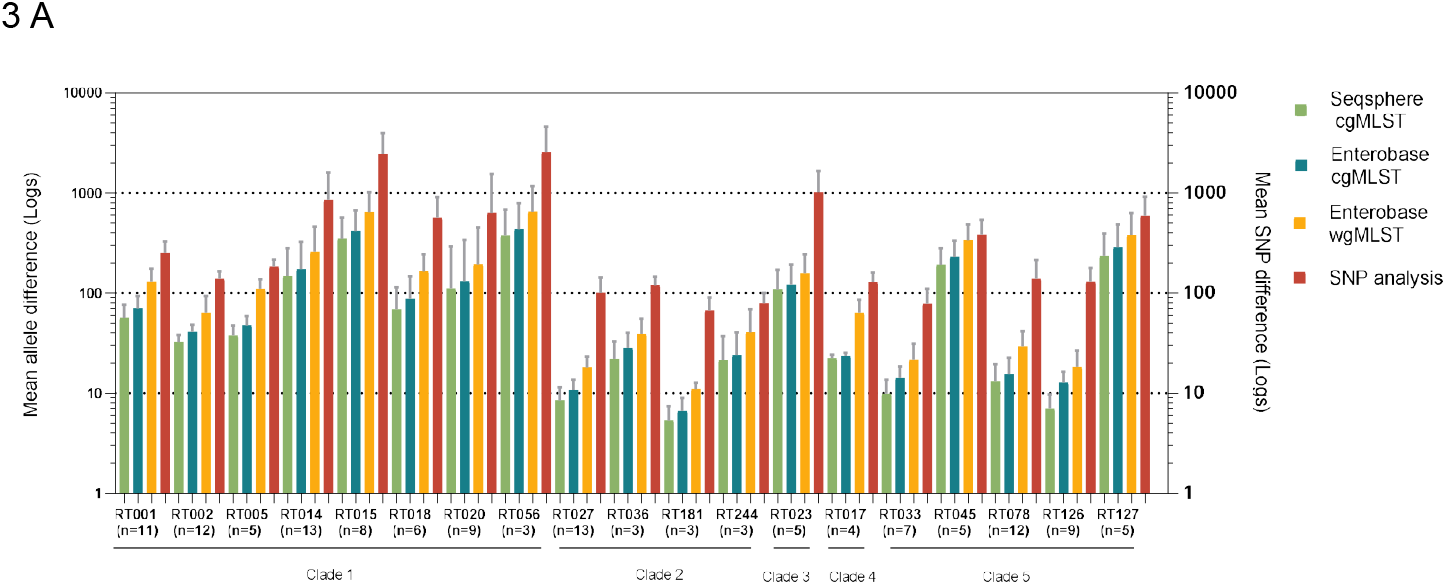
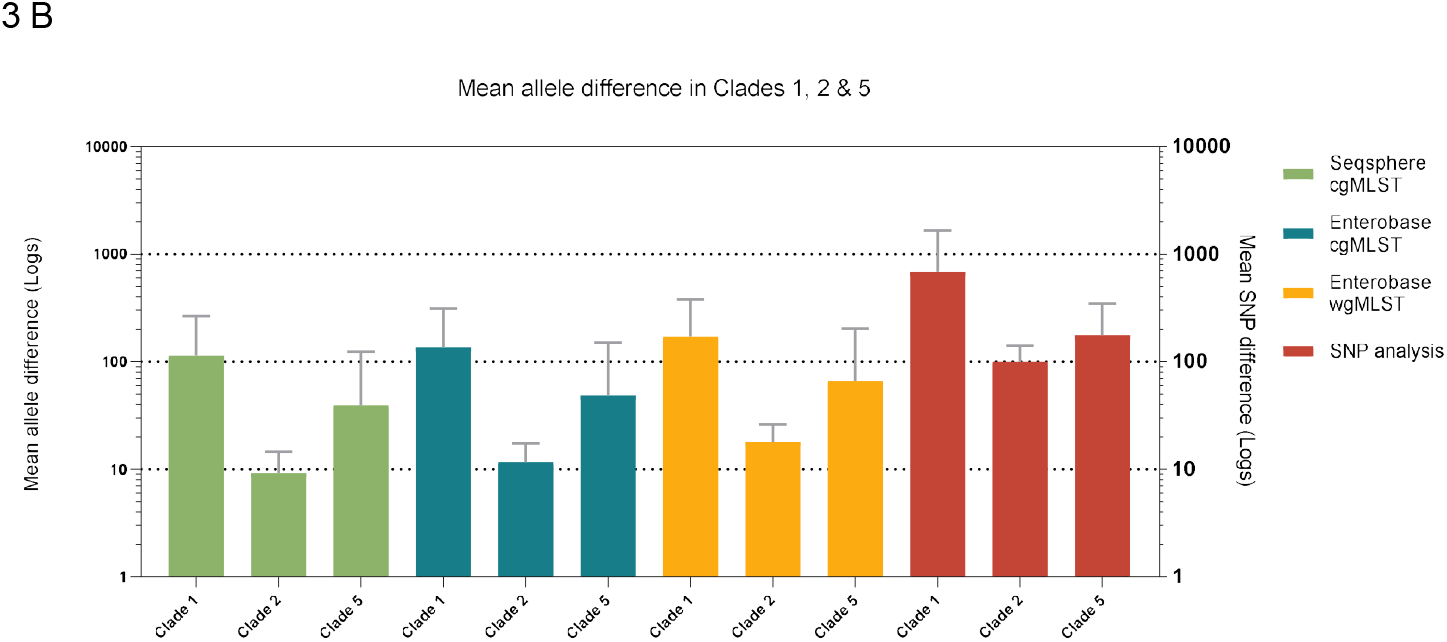
A) Mean intra-ribotype allele and SNP difference shown for ribotypes from MLST Clade 1 (RT001-RT056), Clade 2 (RT027-RT244), Clade 3 (RT023), Clade 4 (RT017) and Clade 5 (RT033-RT127). Mean intra-ribotype allele difference per ribotype is shown in light green, turquois and orange for SqSp cgMLST, EB cgMLST and EB wgMLST, respectively. Mean intra-ribotype SNP difference per ribotype is shown in red. B) Mean intra-ribotype allele and SNP difference shown for MLST Clade 1, Clade 2 and Clade 5. Mean intra-ribotype allele difference per Clade is shown in light green, turquois and orange for SqSp cgMLST, EB cgMLST and EB wgMLST, respectively. Mean intra-ribotype SNP difference per Clade is shown in red.

The mean intra-ribotype allele difference per clade was also calculated for clades 1, 2 and 5 by combining the averages per ribotype within a clade (figure 3B). Clade 1 had the highest average allele difference for SqSp cgMLST, EB cgMLST, EB wgMLST and SNP analysis (114, 136, 171 allele difference and 685 SNPs, respectively). Followed by clade 5 with 39,49, 66 allele differences and 177 SNPs, respectively. Clade 2 had the lowest average intra-ribotype allele difference (9, 12, 18 allele differences and 100 SNPs, respectively).

Clade 1 had the highest mean intra-ribotype allele difference for wgMLST and SNP analysis (171 alleles and 685 SNPs), followed by clade 5 with 66 alleles and 177 SNPs. Clade 2 had again the lowest mean intra-ribotype allele difference (18 alleles and 100 SNPs).

### WGS based typing methods cannot distinguish outbreak strains from non-outbreak strains in ribotypes with a low intra-ribotype allele difference

CE-PCR ribotyping has a low resolution in comparison with whole genome-based typing for outbreak analysis. However, even with the increased resolution of WGS based typing, it remains crucial to understand what defines an outbreak. Bletz *et al*. proposed a threshold of ≤ 6 alleles for cgMLST for isolates that are expected to belong to the same clone (14). In order to guide the interpretation of Bletz *et al*. we compared cgMLST, wgMLST and SNP analysis in 2 suspected outbreak settings. We selected outbreak strains from MLST clades 2 (RT 181) and 5 (RT 078), since both clades have a lower average allele difference. Confirmed outbreak strains were defined as having an epidemiological link (e.g. nursed in the same ward) combined with ≤ 6 allele differences. Control strains belonged to similar PCR ribotypes as the outbreaks strains or to other PCR ribotypes from the same clade.

Next, we analysed the distance matrices of two clusters containing confirmed outbreaks and non-outbreak strains with cgMLST, wgMLST and SNP analysis. The strains within each cluster were either labelled as outbreak strain or control strain. These distance matrices of both clusters were visualized in graphs (Figure 5A and B) with each data point representing a distance in alleles or SNPs between 2 strains. We calculated the range of allele or SNP difference of outbreak strains (Range O) and compared it with the range of allele or SNP difference of non-outbreak strains (Range NO). The area between the upper limit of range O and the lower limit of range NO determines the area where adjustment of the threshold is possible, provided that outbreak strains and non-outbreak strains do not overlap. The larger the area, the better the method can discriminate between outbreak and non-outbreak strains.

The first CDI suspected outbreak we analysed was due to RT078 (clade 5) in a Dutch general hospital, involving 6 patients in the Gastroenterology ward between October-December 2018 (figure 4A). Three patients had an hospital-onset of CDI and 3 had a community-onset (including the index case). The first case (patient A) was admitted on 1^st^ of November and had a community-onset of CDI since the 25^th^ of October. The second case (patient B) was admitted on the 2^nd^ of November and developed hospital-onset of CDI on the 5^th^. The 3^rd^ case (patient C) was admitted on the 12^th^ of November and developed hospital-onset of CDI on the 16^th^. One patient (patient D) was transferred from another hospital on the 24^th^ of November and had CDI since the 13^th^, this patient did not belong to the outbreak. Two other patients (patient E & F) had a community onset of CDI and were admitted both on the 4^th^ of December and had CDI since the 28^th^ of November and 1^st^ of December, respectively. Three isolates from 3 different patients showed a clustering and had 0 allele differences (patients A, B and C), the other 3 patients (patients D, E and F) did not belong to this cluster and had >6 allele differences. Twelve additional control samples from Clade 5 were added to this collection. These included five Leeds-Leiden reference strains (RT033, RT045, RT066, RT078 and RT126) and 7 other strains (RT045, RT066, RT126, RT127 and RT078 (N=3)). Figure 4A depicts the minimum-spanning tree (based on SqSp cgMLST) of the studied isolates of clade 5 (N=18). This resulted in three clusters (≤ 6 alleles), each comprising of epidemiologically related and unrelated strains of which cluster 1 is the largest, involving three strains of the confirmed RT078 outbreak (3 cases [patient A, B and C] and 1 non-case [patient E]) and three control strains (RT066, RT078 and RT126). The second outbreak (18) occurred in a Greek 180-bed rehabilitation clinic involving 15 CDI patients infected with RT181 (clade 2) at the orthopaedics and neurological wards between March and April 2019 (Figure 4B). All 15 patient isolates showed allele differences between 0-2 alleles. Seven control samples from Clade 2 were added to this collection, including Leeds-Leiden reference strains of RT016, RT027, RT198, 1 strain of RT036 and RT176 and 2 strains of RT181. Figure 4B shows the minimum-spanning tree based on SqSp cgMLST. Two clusters could be recognized, each comprising epidemiologically related and unrelated strains. Cluster 1 contained both confirmed outbreak strains (RT181, N=15) and control strains (RT181, N=2). Therefore, the current threshold of ≤6 alleles is not suitable to recognise an outbreak of RT 181.

**Figure 4:**
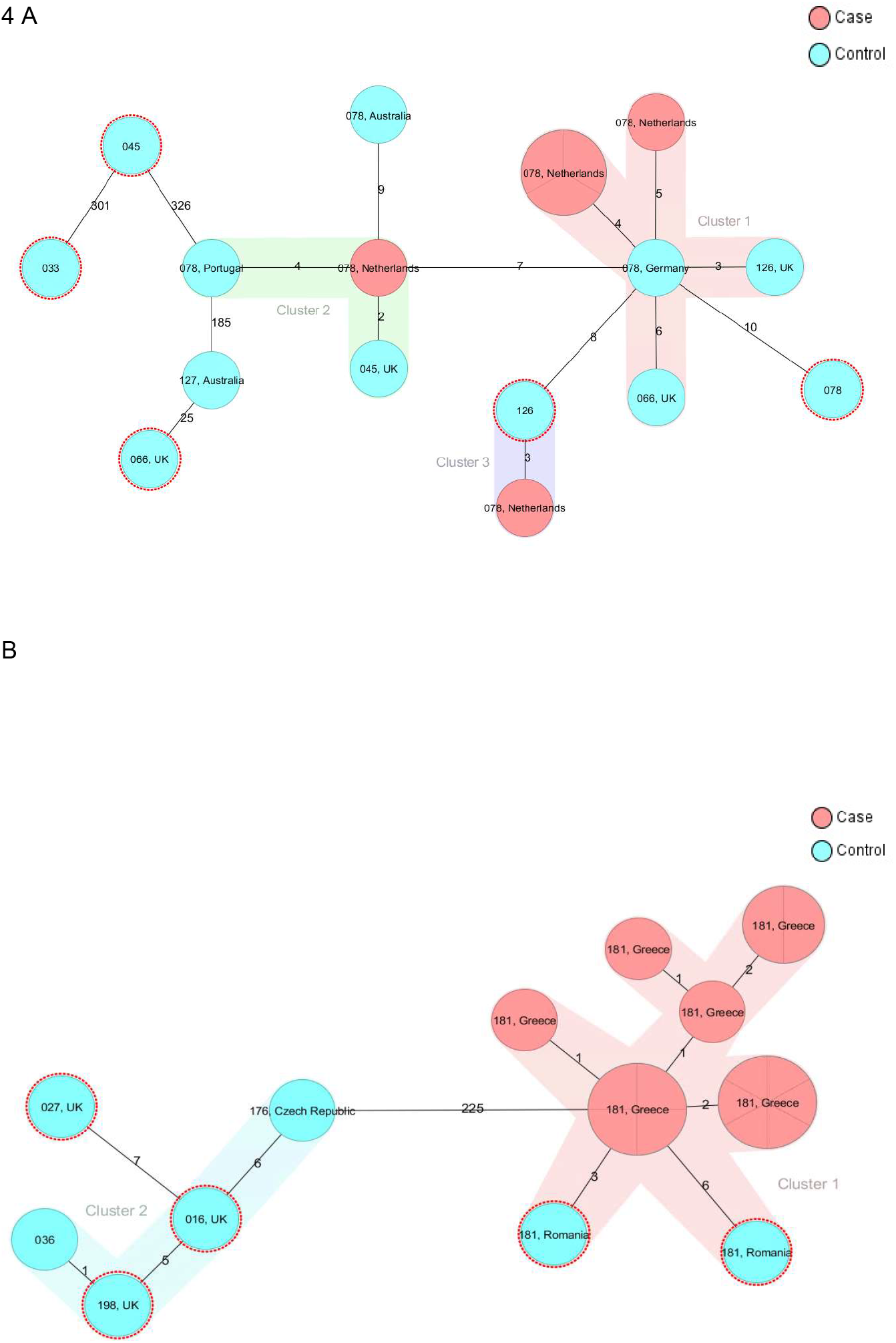
SqSP cgMLST analysis with minimum-spanning trees of 2 suspected CDI outbreaks of RT078 and RT181. A) Minimum-spanning tree of PCR-ribotype 078 (clade 5) CDI suspected outbreak with 6 cases (RT078, shown in red), confirmed outbreak with 3 cases (RT078, shown in largest red circle) and added control strains of ribotypes belonging to clade 5 (reference strains of RT033, RT045, RT066, RT078, RT126 shown in blue with red circles and non-reference strains of RT045, RT066, RT078, RT126 and RT127 shown in blue). B) Minimum-spanning tree of PCR-ribotype 181 (clade 2) CDI suspected outbreak with 15 suspected and confirmed cases (RT181, shown in red) and control strains of ribotypes of clade 2 (reference strains of RT016, RT027, RT181 and RT198 shown in blue with red circles and non-reference strains of RT036 and RT176 shown in blue. The size and septation of the circle in the minimum-spanning trees corresponds to the number of included strains. The numbers between each circle correspond to the number of different alleles between the strains. The coloured shadowing of circles represents a cluster with <= 6 allele differences that are genetically related. One or more strains inside a circle means that these strains have 0 allele difference.

Figure 5A shows that all WGS method could distinguish between confirmed outbreak and non-outbreak RT 078 strains, since there is no overlap between range O and range NO. SNP analysis had the best discriminatory power, followed by EB wgMLST and cgMLST, which showed the lowest discriminatory power. Figure 5B shows that wgMLST is the only method that could discriminate between outbreak and non-outbreak RT 181 strains, whereas cgMLST and SNP analysis show overlap in their ranges. Ranges O and NO are shown in Table 2 for both clusters and all applied typing methods. No overlap was seen between Range O and Range NO from Cluster 1 from the RT078 CDI outbreak. For SqSp cgMLST and EB cgMLST cluster 1 showed a difference of 3 alleles and 2 alleles between the Range O and Range NO, respectively. Furthermore, the difference between Range O and Range NO was for wgMLST and SNP analysis 6 alleles and 8 SNPs, indicating that the threshold could be lowered. However, Cluster 1 from the RT181 CDI outbreak showed overlap between Range O and Range NO in cgMLST and SNP analysis, but not in wgMLST, suggesting that the threshold only could be adjusted in wgMLST.

**Figure 5:**
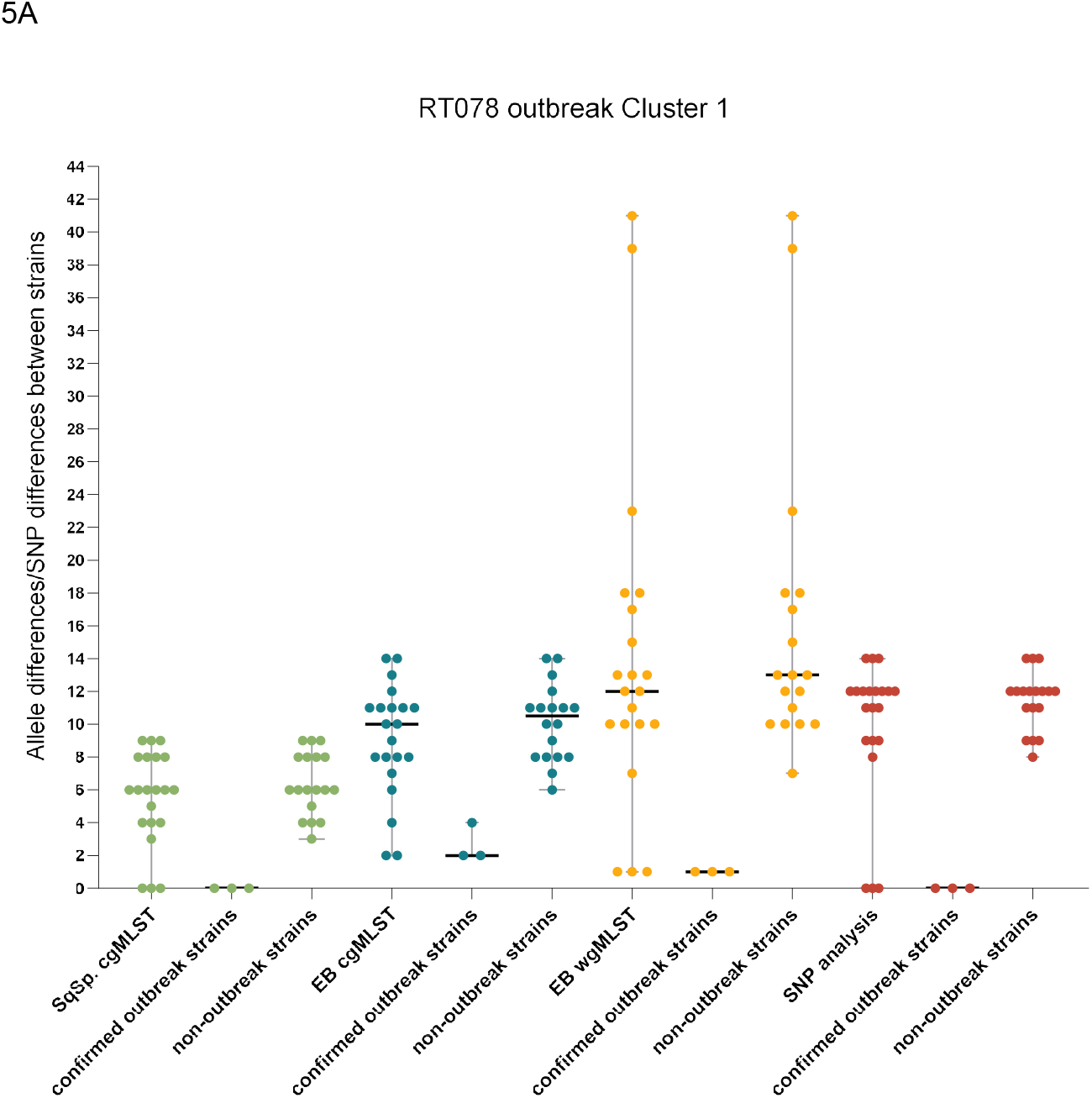
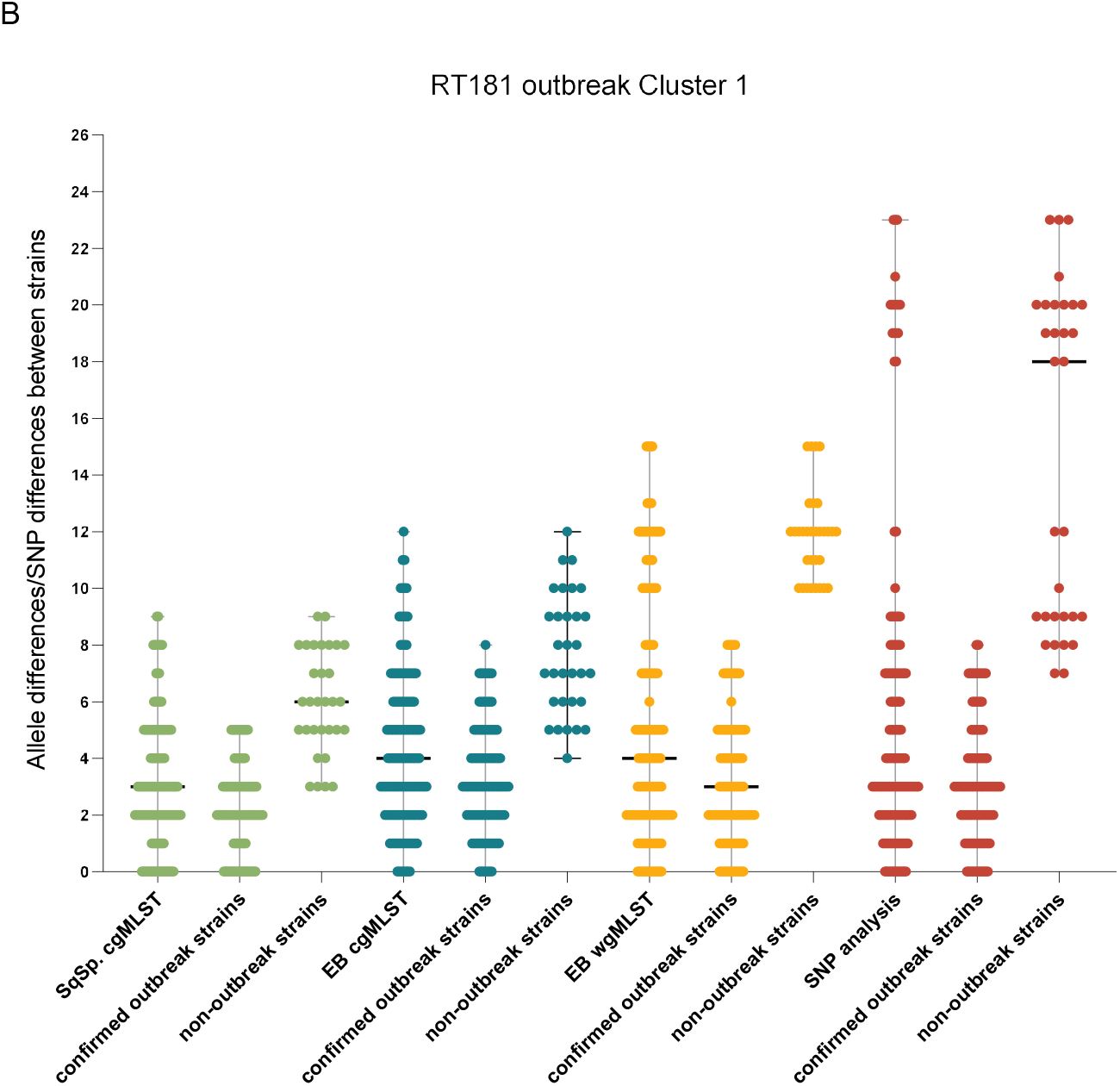
Visualised distance matrices of strain pairs based on cgMLST, wgMLST and SNP analysis of isolates of cluster 1 as described in figure 4A & 4B. A) Visualised distance matrix of strain pairs belonging to cluster 1 of RT078. B) Visualised distance matrix of strain pairs belonging to cluster 1 of RT181. Allele difference per pair of strains is shown in light green, turquois and orange for cgMLST in SeqSphere, cgMLST and wgMLST in EnteroBase, respectively. SNP difference per pair of strains is shown in red.

**Table 2:** comparison in range between outbreak and non-outbreak strains of RT078 and RT181.

| Typing method | Strains | Range (alleles or SNPs) | Difference between Range O & Range NO <sup>a</sup> |
| --- | --- | --- | --- |
| SqSp cgMLST | 078 confirmed outbreak | 0 | 3 |
|  | non-outbreak | 3-9 |  |
|  | 181 confirmed outbreak | 0-5 | overlap |
|  | non-outbreak | 3-9 |  |
| EB cgMLST | 078 confirmed outbreak | 2-4 | 2 |
|  | non-outbreak | 6-14 |  |
|  | 181 confirmed outbreak | 0-8 | overlap |
|  | non-outbreak | 4-12 |  |
| EB wgMLST | 078 confirmed outbreak | 1 | 6 |
|  | non-outbreak | 7-41 |  |
|  | 181 confirmed outbreak | 0-8 | 2 |
|  | non-outbreak | 10-15 |  |
| SNP analysis | 078 confirmed outbreak | 0 | 8 |
|  | non-outbreak | 8-14 |  |
|  | 181 confirmed outbreak | 0-9 | overlap |
|  | non-outbreak | 7-23 |  |

a) Range O is the range in allele or SNP difference between all outbreak strains. Range NO is the range in allele or SNP difference of non-outbreak strains compared with themselves and compared with outbreak strains.

## DISCUSSION

We tested the backward compatibility between SqSp cgMLST and CE-PCR ribotyping and found 82 of 100 different PCR ribotypes had a unique cgMLST profile using a cut-off of ≤6 alleles differences. Assessing the performance of cgMLST, wgMLST and SNP typing in comparison with CE-PCR ribotyping revealed that intra-ribotype alleles difference varied per ribotype and per MLST clade. Application of cg/wgMLST and SNP analysis in outbreak settings of RT078 and RT181 showed that these methods can only distinguish outbreak strains from non-outbreak strains when a cut-off threshold of 3 alleles is used.

We show that SqSp cgMLST is backward compatible with CE-PCR-ribotyping, but there are certain ribotypes that are indistinguishable by SqSp cgMLST. These data are consistent with Seth-Smith *et al*. who found different PCR ribotypes (RT 078-126, RT 106-RT500) clustering with maximum of 9 allelic difference. In agreement with the findings of Seth-Smith et al., we found ribotypes from clade 2 and 5 with the lowest mean intra-ribotype allele difference. We applied in our study the average allele differences, contrary to the study of Seth-Smith who used the maximum allelic difference. When we analysed for the maximum allelic difference, we found higher differences in all studied ribotypes than Seth-Smith *et al*. (e.g. RT027: 12 allelic difference vs. 16 allelic difference in our study; RT078: 9 vs. 28, respectively; RT023: 52 vs. 199, respectively). This may have been caused by the selection of samples, since we excluded samples from outbreaks by selecting strains separated in time and space.

Our results are also consistent with another study (30) that used SNP analysis to investigate the diversity within a ribotype. The study showed that MLST ST1 (correlates with ribotype 027) was genetically less diverse with a lower SNP distance range between isolates than ST2 (correlates with ribotype 014). Finally, Frentrup *et al*, observed clustering of several ribotypes (e.g. RT001/RT241, RT106/RT500 and RT078/RT126) from MLST clades 1 and 5 (16), also in agreement with our observations.

Interestingly, decreasing the threshold from 6 to 0 allele difference will still result in clustering of certain ribotypes. The clustering between two strains of RT045 and two strains of RT127 at a threshold of 0 alleles in SqSp cgMLST was verified with EB cgMLST and SNP analysis. With EB cgMLST one clustering pair of RT045 and RT127 showed 1 allele difference, whereas the other remained at 0 allele difference. Verification with SNP analysis showed 2 and 7 SNP differences. This observation impairs the backward compatibility of cgMLST with CE-PCR ribotyping and excludes studying the epidemiological links of some strains belonging to RT045 and RT127. Our results demonstrate that the mean allele differences between strains from the same PCR ribotype with SqSp cgMLST and EB cgMLST are lower in comparison with EB wgMLST and SNP analysis, with the latter showing the highest resolution. Similar results were seen in the studied RT078 CDI outbreak, where EB wgMLST and SNP analysis showed more discriminatory power in comparison with cgMLST. Interestingly, EB wgMLST was the only WGS based method that could discriminate between outbreak strains and non-outbreak strains in RT181 CDI outbreak. A reason could be that EB wgMLST uses a pangenome as a scheme consisting of several *C. difficile* genomes, in contrast with SNP analysis, which used strain 630 as the reference genome. Ribotypes from clades (e.g. clade 2) that have emerged relatively recently will have lower mean intra-ribotype allele differences as strains from these ribotypes look genetically more similar. Therefore, it may be challenging to distinguish which strains are involved in an outbreak. Another problem with these recently emerged ribotypes (e.g. RT181) is that we have limited data to assess the intra-ribotype allele difference more accurately.

Based on our observations in two CDI outbreaks, we conclude that WGS based methods cannot discriminate between outbreak and non-outbreak strains in MLST clades with low intra-ribotype allele difference. It remains unknown why some clades are less diverse. It is possible that they have emerged relatively recently and therefore are less diverse. Alternatively, the strains in these clades could have a lower mutation rate resulting in less diversity and therefore a lower intra-ribotype allele difference (31), (32). For outbreaks caused by PCR ribotypes belonging to other clades than 2 and 5, the performance of cgMLST is comparable with SNP analysis. Our results are consistent with other studies showing a comparable performance of cgMLST with SNP analysis (14), (33). Based upon the Oxfordshire data set (31), Frentrup *et al*. had a similar conclusion regarding cgMLST and SNP analysis (16). They showed that *C. difficile* genomes that differ by ≤2 alleles generally also differ by 2 ≤SNPs, using a logistic regression model, and concluded that cgMLST is equivalent to SNP analysis for identifying transmission chains between patients. Bletz *et al*. showed similar results between cgMLST and SNP analysis in detecting clusters when an outbreak due to ST1 was investigated (14). The main strength of our study is that we compared the performance of several typing methods, in contrast to previous studies (14), (11), (15), (16). We also expanded the collection of *C. difficile* strains and tested more than 600 sequenced strains belonging to 100 unique ribotypes. Our study has also some limitations. The lack of sufficient available genome sequences from strains belonging to clades 3 and 4 limits the generalizability of our findings. Though the backward compatibility was not tested for EB wgMLST, the results can be extrapolated from SqSp cgMLST, EB cgMLST and SNP analysis, since the discriminatory power of EB wgMLST lies between the latter two. We could not verify the correctness of the strain PCR ribotypes, as we had only access to the information as deposited by the researchers. There are a few ribotypes that have similar banding pattern and could be misidentified. The best example is the similarity of RT014 with RT020; they have an almost identical PCR banding pattern, but they differ substantially from each other by cgMLST. Though we only studied two outbreaks, we carefully selected the outbreaks by choosing PCR ribotypes with low intra-ribotype alleles variation. Finally, we have not tested long read sequencing from which theoretically in silico PCR ribotyping can also be obtained.

We propose to decrease the current threshold of 6 alleles (14) to 3 alleles when using cgMLST in outbreak situations. We found a difference of 2 and 3 alleles between controls and outbreak strains with EB cgMLST and SqSp cgMLST, respectively. In the study by Eyre *et al*. the evolutionary rate of *C. difficile* was estimated to be 0.74 SNVs (95% confidence interval, 0.22-1.40) per genome per year (34). They expected 0-2 SNPs to occur when isolates are obtained <124 days apart and 3 SNPs when isolates were obtained 124-364 days apart. However, only vegetative *C. difficile* isolates obtained from patients were analyzed. According to Weller & Wu sporulation reduces the evolutionary rate of *Firmicutes* (35). Therefore, we expect that the evolutionary rate of *C. difficile* is lower during CDI transmission than during CDI within a patient, since the spores need time to transmit to another patient and otherwise lie dormant in the surroundings in a healthcare facility or in the environment for a long period. Accordingly, we expect that outbreak strains will generally fall within 0-2 alleles.

Nevertheless, we recommend a threshold of 3 alleles to compensate for any assembly artifacts when less conservative pipelines are used and for outbreaks that last longer than 124 days (36).

A concern with application of cgMLST is the availability of various cgMLST schemes and software programs. The centralized databases need resources to maintain their databases of sequentially numbered alleles. To tackle the problem of the need for a centralized database and to rapidly identify related genomes against a background of thousands of other identified genomes, Hash-Based cgMLST has been developed (11). It is based on cgMLST, but converts alleles in a unique hash or short string of letters. Furthermore, if every software provider uses its own cgMLST scheme, inter-laboratory comparison is delayed and understanding of epidemiology is hampered. As Werner *et al*. proposed, it is favourable that a fixed cgMLST scheme is constructed (33). Furthermore, there are logistical and cost considerations for routine implementation of cgMLST. Reference laboratory are needed with a good infrastructure to sequence strains on a routine basis while keeping the costs in mind as well.

In summary, cgMLST has the potential to replace CE-PCR ribotyping for *C. difficile*. The method provides similar differentiation of strains, is easy to standardize, is reproducible and shows a high discriminatory power. Several cgMLST based typing methods have emerged with all their specific advantages and disadvantages (11), (14), (16). For the time being, it remains unclear whether one method will get the preference over other methods or that every center will use its own method. However, it is important to ensure that local and international strains can be compared regardless of the use of different methods either by exchange of raw data or via a centralized multi-national database with a fixed cgMLST scheme where every center contributes to. A consensus group could be assembled to harmonize these efforts as has been done previously for CE-PCR ribotyping (4).

## Acknowledgments

We thank B.V.H. Hornung for his assistance with data collection.

---

**Table S1:** included WGS strains of *C. difficile* in this study.

| Sample ID | Ribotype | Collection Date | Country of Isolation | ST | CC | SRA accession no |
| --- | --- | --- | --- | --- | --- | --- |
| SRR7308630-001 | 001 | 2013 | UK | 3 | 1 | SRR7308630 |
| SRR1519369-LL-001 | 001 | ? | ? | 3 | 1 | SRR1519369 |
| SRR7309226-001 | 001 | 2013 | Netherlands | 3 | 1 | SRR7309226 |
| SRR7308692-001 | 001 | 2013 | Italy | 3 | 1 | SRR7308692 |
| SRR7308732-001 | 001 | 2013 | Finland | 3 | 1 | SRR7308732 |
| SRR7308761-001 | 001 | 2013 | Sweden | 3 | 1 | SRR7308761 |
| SRR7308773-001 | 001 | 2013 | Slovakia | 3 | 1 | SRR7308773 |
| SRR7308776-001 | 001 | 2013 | Slovakia | 3 | 1 | SRR7308776 |
| SRR7308804-001 | 001 | 2013 | Spain | 3 | 1 | SRR7308804 |
| SRR7308833-001 | 001 | 2013 | France | 3 | 1 | SRR7308833 |
| SRR7308981-001 | 001 | 2013 | Italy | 3 | 1 | SRR7308981 |
| SRR7309099-001 | 001 | 2013 | Germany | 3 | 1 | SRR7309099 |
| SRR7309176-001 | 001 | ? | Bulgaria | 3 | 1 | SRR7309176 |
| SRR7308836-001 | 001 | ? | Netherlands | 3 | 1 | SRR7308836 |
| SRR7308645-002 | 002 | 2013 | Portugal | 8 | 1 | SRR7308645 |
| SRR7308677-002 | 002 | 2013 | Netherlands | 8 | 1 | SRR7308677 |
| SRR7309212-002 | 002 | 2013 | Belgium | 8 | 1 | SRR7309212 |
| SRR7308752-002 | 002 | 2013 | France | 8 | 1 | SRR7308752 |
| SRR7309014-002 | 002 | 2013 | Sweden | 8 | 1 | SRR7309014 |
| SRR7309032-002 | 002 | 2013 | Finland | 8 | 1 | SRR7309032 |
| SRR7309128-002 | 002 | 2013 | Romania | 8 | 1 | SRR7309128 |
| SRR7309141-002 | 002 | 2013 | Portugal | 8 | 1 | SRR7309141 |
| SRR7309152-002 | 002 | 2013 | Romania | 8 | 1 | SRR7309152 |
| SRR7309217-002 | 002 | 2013 | Italy | 8 | 1 | SRR7309217 |
| SRR7309219-002 | 002 | 2013 | UK | 8 | 1 | SRR7309219 |
| SRR1519370-LL-002 | 002 | ? | ? | 8 | 1 | SRR1519370 |
| SRR6042346-002 | 002 | 2010 | USA | 55 | 1 | SRR6042346 |
| SRR7308785-002 | 002 | 2013 | Poland | 8 | 1 | SRR7308785 |
| SRR7308659-002 | 002 | 2013 | Germany | 8 | 1 | SRR7308659 |
| SRR7308698-002 | 002 | 2013 | UK | 8 | 1 | SRR7308698 |
| SRR7308733-002 | 002 | 2013 | Italy | 8 | 1 | SRR7308733 |
| SRR7308704-002 | 002 | 2013 | France | 8 | 1 | SRR7308704 |
| SRR1519371-LL-003 | 003 | 2005 | UK | 12 | 1 | SRR1519371 |
| SRR7852176-003 | 003 | ? | UK | 12 | 1 | SRR7852176 |
| SRR1519372-LL-004 | 004 | 1995 | UK | 115 | 1 | SRR1519372 |
| SRR7852181-005 | 005 | ? | UK | 6 | 1 | SRR7852181 |
| SRR7852186-005 | 005 | ? | UK | 6 | 1 | SRR7852186 |
| SRR7852185-005 | 005 | ? | UK | ? | ? | SRR7852185 |
| SRR6042365-005 | 005 | 2010 | USA | 6 | 1 | SRR6042365 |
| ERR833662-005 | 005 | ? | UK | 6 | 1 | ERR833662 |
| SRR6042356-005 | 005 | 2010 | USA | 6 | 1 | SRR6042356 |
| SRR7852187-005 | 005 | ? | UK | 6 | 1 | SRR7852187 |
| SRR7852208-005 | 005 | ? | UK | 6 | 1 | SRR7852208 |
| SRR6042370-005 | 005 | 2010 | USA | 6 | 1 | SRR6042370 |
| SRR1519373-LL-005 | 005 | 2005 | UK | 6 | 1 | SRR1519373 |
| SRR1519374-LL-006 | 006 | 1995 | UK | 2 | 1 | SRR1519374 |
| SRR1519375-LL-007 | 007 | 1995 | UK | 49 | 1 | SRR1519375 |
| SRR7852197-007 | 007 | ? | UK | 49 | 1 | SRR7852197 |
| SRR7852191-009 | 009 | ? | UK | 3 | 1 | SRR7852191 |
| SRR1519376-LL-009 | 009 | 2007 | UK | 3 | 1 | SRR1519376 |
| SRR1519377-LL-010 | 010 | ? | UK | 15 | 1 | SRR1519377 |
| ERR833666-010 | 010 | ? | UK | 15 | 1 | ERR833666 |
| ERR833672-010 | 010 | ? | UK | 15 | 1 | ERR833672 |
| SRR1519378-LL-011 | 011 | 2005 | UK | 36 | 1 | SRR1519378 |
| ERR833660-012 | 012 | ? | UK | 54 | 1 | ERR833660 |
| SRR1519379-LL-012 | 012 | ? | Belgium | 54 | 1 | SRR1519379 |
| SRR593175-013 | 013 | ? | USA | 45 | 1 | SRR593175 |
| ERR1307016-014 | 014 | ? | Australia | 2 | 1 | ERR1307016 |
| SRR7308710-014 | 014 | 2013 | Hungary | 13 | 1 | SRR7308710 |
| SRR7308735-014 | 014 | 2013 | Sweden | 49 | 1 | SRR7308735 |
| SRR7308765-014 | 014 | 2013 | Czech Republic | 2 | 1 | SRR7308765 |
| SRR7308801-014 | 014 | 2013 | Italy | 2 | 1 | SRR7308801 |
| SRR7309218-014 | 014 | 2013 | Ireland | 2 | 1 | SRR7309218 |
| SRR7308852-014 | 014 | 2013 | UK | 13 | 1 | SRR7308852 |
| SRR7308907-014 | 014 | 2013 | Italy | 2 | 1 | SRR7308907 |
| SRR7308925-014 | 014 | 2013 | Czech Republic | 2 | 1 | SRR7308925 |
| SRR7308936-014 | 014 | 2013 | Slovakia | 13 | 1 | SRR7308936 |
| SRR7309040-014 | 014 | 2013 | Spain | 2 | 1 | SRR7309040 |
| SRR7309087-014 | 014 | 2013 | Romania | ? (unknown) | ? | SRR7309087 |
| SRR7309095-014 | 014 | 2013 | Sweden | ? (unknown) | ? | SRR7309095 |
| SRR7309127-014 | 014 | 2013 | Germany | 2 | 1 | SRR7309127 |
| SRR7309213-014 | 014 | 2013 | UK | 14 | 1 | SRR7309213 |
| SRR7308784-014 | 014 | 2013 | Hungary | 2 | 1 | SRR7308784 |
| SRR1519380-LL-014 | 014 | ? | ? | 2 | 1 | SRR1519380 |
| SRR7309104-014 | 014 | 2013 | France | 2 | 1 | SRR7309104 |
| SRR7309188-014 | 014 | 2013 | Poland | 2 | 1 | SRR7309188 |
| SRR7308861-014 | 014 | 2013 | UK | 2 | 1 | SRR7308861 |
| SRR7309062-015 | 015 | 2013 | UK | 44 | 1 | SRR7309062 |
| SRR7309143-015 | 015 | 2013 | UK | 44 | 1 | SRR7309143 |
| SRR7309211-015 | 015 | 2013 | Germany | 10 | 1 | SRR7309211 |
| SRR7308686-015 | 015 | 2013 | Germany | 10 | 1 | SRR7308686 |
| SRR7308700-015 | 015 | 2013 | Germany | 44 | 1 | SRR7308700 |
| SRR7308914-015 | 015 | 2013 | Germany | 44 | 1 | SRR7308914 |
| SRR7308766-015 | 015 | 2013 | Bulgaria | 44 | 1 | SRR7308766 |
| SRR7308811-015 | 015 | 2013 | Netherlands | 160 | 1 | SRR7308811 |
| SRR7308858-015 | 015 | 2013 | Germany | 10 | 1 | SRR7308858 |
| SRR7308881-015 | 015 | 2013 | France | 10 | 1 | SRR7308881 |
| SRR7308931-015 | 015 | 2013 | Bulgaria | 44 | 1 | SRR7308931 |
| SRR7309068-015 | 015 | 2013 | UK | 44 | 1 | SRR7309068 |
| SRR7309165-015 | 015 | 2012 | France | 10 | 1 | SRR7309165 |
| SRR1519381-LL-015 | 015 | 2006 | UK | 10 | 1 | SRR1519381 |
| SRR7308670-015 | 015 | 2013 | Romania | 44 | 1 | SRR7308670 |
| SRR7308847-015 | 015 | 2013 | Ireland | 44 | 1 | SRR7308847 |
| SRR7308958-015 | 015 | 2013 | UK | 44 | 1 | SRR7308958 |
| SRR7308789-015 | 015 | 2012 | France | 10 | 1 | SRR7308789 |
| SRR7309191-015 | 015 | 2013 | Germany | 10 | 1 | SRR7309191 |
| SRR6042362-015/046 | 015/046 | 2010 | USA | 10 | 1 | SRR6042362 |
| SRR1519382-LL-016 | 016 | 2008 | UK | 1 | 2 | SRR1519382 |
| ERR1346261-017 | 017 | ? | ? | 37 | 4 | ERR1346261 |
| ERR1346244-017 | 017 | ? | ? | 37 | 4 | ERR1346244 |
| ERR788991-017 | 017 | ? | UK | 37 | 4 | ERR788991 |
| ERR1346256-017 | 017 | ? | ? | 37 | 4 | ERR1346256 |
| ERR1346252-017 | 017 | ? | ? | 37 | 4 | ERR1346252 |
| ERR1346250-017 | 017 | ? | ? | 37 | 4 | ERR1346250 |
| ERR1346249-017 | 017 | ? | ? | 37 | 4 | ERR1346249 |
| ERR1346248-017 | 017 | ? | ? | 37 | 4 | ERR1346248 |
| ERR1346246-017 | 017 | ? | ? | 37 | 4 | ERR1346246 |
| ERR1346245-017 | 017 | ? | ? | 37 | 4 | ERR1346245 |
| ERR1346263-017 | 017 | ? | ? | 37 | 4 | ERR1346263 |
| ERR1346247-017 | 017 | ? | ? | 37 | 4 | ERR1346247 |
| ERR1346253-017 | 017 | ? | ? | 37 | 4 | ERR1346253 |
| ERR1346257-017 | 017 | ? | ? | 37 | 4 | ERR1346257 |
| ERR1346255-017 | 017 | ? | ? | 37 | 4 | ERR1346255 |
| ERR788960-017 | 017 | ? | UK | 37 | 4 | ERR788960 |
| ERR788962-017 | 017 | ? | UK | 37 | 4 | ERR788962 |
| ERR788967-017 | 017 | ? | UK | 37 | 4 | ERR788967 |
| ERR788968-017 | 017 | ? | UK | 37 | 4 | ERR788968 |
| ERR788969-017 | 017 | ? | UK | 37 | 4 | ERR788969 |
| ERR788971-017 | 017 | ? | UK | 37 | 4 | ERR788971 |
| ERR788972-017 | 017 | ? | UK | 37 | 4 | ERR788972 |
| ERR788974-017 | 017 | ? | UK | 37 | 4 | ERR788974 |
| ERR788975-017 | 017 | ? | UK | 37 | 4 | ERR788975 |
| ERR788976-017 | 017 | ? | UK | 37 | 4 | ERR788976 |
| ERR788977-017 | 017 | ? | UK | 37 | 4 | ERR788977 |
| ERR788978-017 | 017 | ? | UK | 37 | 4 | ERR788978 |
| ERR788979-017 | 017 | ? | UK | 37 | 4 | ERR788979 |
| ERR788980-017 | 017 | ? | UK | 37 | 4 | ERR788980 |
| ERR788981-017 | 017 | ? | UK | 37 | 4 | ERR788981 |
| ERR1346259-017 | 017 | ? | ? | 37 | 4 | ERR1346259 |
| ERR1346258-017 | 017 | ? | ? | 37 | 4 | ERR1346258 |
| ERR1346262-017 | 017 | ? | ? | 37 | 4 | ERR1346262 |
| ERR1346260-017 | 017 | ? | ? | 37 | 4 | ERR1346260 |
| ERR788994-017 | 017 | ? | UK | 37 | 4 | ERR788994 |
| ERR788984-017 | 017 | ? | UK | 37 | 4 | ERR788984 |
| ERR1346251-017 | 017 | ? | ? | 37 | 4 | ERR1346251 |
| ERR788985-017 | 017 | ? | UK | 37 | 4 | ERR788985 |
| ERR788986-017 | 017 | ? | UK | 37 | 4 | ERR788986 |
| ERR788987-017 | 017 | ? | UK | 37 | 4 | ERR788987 |
| ERR788989-017 | 017 | ? | UK | 37 | 4 | ERR788989 |
| ERR788993-017 | 017 | ? | UK | 37 | 4 | ERR788993 |
| ERR788995-017 | 017 | ? | UK | 37 | 4 | ERR788995 |
| SRR1519383-LL-017 | 017 | ? | ? | 37 | 4 | SRR1519383 |
| SRR7309227-018 | 018 | 2013 | Italy | 17 | 1 | SRR7309227 |
| SRR7308628-018 | 018 | 2013 | Italy | 17 | 1 | SRR7308628 |
| SRR7308794-018 | 018 | 2013 | Italy | 17 | 1 | SRR7308794 |
| SRR7309012-018 | 018 | 2013 | Italy | 17 | 1 | SRR7309012 |
| SRR7309013-018 | 018 | 2013 | Italy | 17 | 1 | SRR7309013 |
| SRR7309172-018 | 018 | 2013 | Italy | 17 | 1 | SRR7309172 |
| SRR7308763-018 | 018 | 2013 | Italy | 17 | 1 | SRR7308763 |
| SRR7309195-018 | 018 | 2013 | Italy | 17 | 1 | SRR7309195 |
| SRR7308787-018 | 018 | 2013 | Italy | 17 | 1 | SRR7308787 |
| SRR7308816-018 | 018 | 2013 | Italy | 17 | 1 | SRR7308816 |
| SRR7308908-018 | 018 | 2013 | Italy | 17 | 1 | SRR7308908 |
| SRR7308901-018 | 018 | 2012 | France | 17 | 1 | SRR7308901 |
| SRR7309096-018 | 018 | 2013 | Germany | 17 | 1 | SRR7309096 |
| SRR7308930-018 | 018 | 2013 | Romania | 17 | 1 | SRR7308930 |
| SRR7309150-018 | 018 | 2013 | Romania | 17 | 1 | SRR7309150 |
| SRR7309190-018 | 018 | 2013 | Poland | 17 | 1 | SRR7309190 |
| SRR1519384-LL-018 | 018 | ? | ? | 17 | 1 | SRR1519384 |
| SRR7309216-018 | 018 | 2013 | Romania | 17 | 1 | SRR7309216 |
| SRR3547038-018/052 | 018/052 | 2015 | Italy | 17 | 1 | SRR3547038 |
| SRR1519385-LL-019 | 019 | ? | ? | 67 | 2 | SRR1519385 |
| SRR7309161-020 | 020 | 2013 | Germany | 2 | 1 | SRR7309161 |
| SRR7308878-020 | 020 | 2012 | France | 2 | 1 | SRR7308878 |
| SRR7308905-020 | 020 | 2013 | Portugal | 2 | 1 | SRR7308905 |
| SRR7308629-020 | 020 | 2012 | France | 2 | 1 | SRR7308629 |
| SRR7308750-020 | 020 | 2013 | Czech Republic | 110 | 1 | SRR7308750 |
| SRR7309214-020 | 020 | 2013 | Italy | 2 | 1 | SRR7309214 |
| SRR7309116-020 | 020 | 2013 | Belgium | 2 | 1 | SRR7309116 |
| SRR7308953-020 | 020 | 2013 | Belgium | 2 | 1 | SRR7308953 |
| SRR7308927-020 | 020 | 2013 | Czech Republic | 2 | 1 | SRR7308927 |
| SRR7308894-020 | 020 | 2013 | France | 2 | 1 | SRR7308894 |
| SRR7308873-020 | 020 | 2013 | Portugal | 2 | 1 | SRR7308873 |
| SRR7308778-020 | 020 | 2013 | Belgium | 2 | 1 | SRR7308778 |
| SRR7308770-020 | 020 | 2013 | Germany | 2 | 1 | SRR7308770 |
| SRR7308759-020 | 020 | 2013 | Romania | 2 | 1 | SRR7308759 |
| SRR7308709-020 | 020 | 2013 | Romania | 2 | 1 | SRR7308709 |
| SRR7308689-020 | 020 | 2013 | Germany | 2 | 1 | SRR7308689 |
| SRR7308688-020 | 020 | 2013 | France | 2 | 1 | SRR7308688 |
| SRR7308644-020 | 020 | 2013 | Italy | 2 | 1 | SRR7308644 |
| SRR1519386-LL-020 | 020 | 1996 | Poland | 13 | 1 | SRR1519386 |
| SRR7308693-020 | 020 | 2013 | Czech Republic | 2 | 1 | SRR7308693 |
| SRR7308969-020 | 020 | 2013 | Germany | 2 | 1 | SRR7308969 |
| SRR1519387-LL-022 | 022 | 1990 | UK | 66 | 1 | SRR1519387 |
| ERR232375-023 | 023 | 2008 | UK | 5 | 3 | ERR232375 |
| ERR2215969-023 | 023 | ? | ? | 5 | 3 | ERR2215969 |
| SRR1519388-LL-023 | 023 | 2001 | Netherlands | 5 | 3 | SRR1519388 |
| ERR125913-Sang-023 | 023 | 2008 | UK | 5 | 3 | ERR125913 |
| ERR142362-023 | 023 | 2007 | UK | 25 | 3 | ERR142362 |
| ERR126268-023 | 023 | 2007 | UK | 22 | 3 | ERR126268 |
| SRR1519389-LL-025 | 025 | 1994 | Ireland | 49 | 1 | SRR1519389 |
| SRR7852202-026 | 026 | ? | UK | 7 | 1 | SRR7852202 |
| SRR7852178-026 | 026 | ? | UK | 7 | 1 | SRR7852178 |
| SRR7852201-026 | 026 | ? | UK | 7 | 1 | SRR7852201 |
| SRR1519390-LL-026 | 026 | 1996 | UK | 7 | 1 | SRR1519390 |
| ERR467623-027 | 027 | ? | ? | 154 | 2 | ERR467623 |
| ERR044837-027 | 027 | 2006 | Netherlands | 1 | 2 | ERR044837 |
| ERR044840-027 | 027 | 2006 | Netherlands | 1 | 2 | ERR044840 |
| ERR044825-027 | 027 | 2006 | Austria | 1 | 2 | ERR044825 |
| ERR044843-027 | 027 | 2006 | Netherlands | 1 | 2 | ERR044843 |
| ERR044846-027 | 027 | 2009 | Poland | 1 | 2 | ERR044846 |
| ERR030295-027 | 027 | 2010 | UK | 1 | 2 | ERR030295 |
| ERR044842-027 | 027 | 2006 | Netherlands | 1 | 2 | ERR044842 |
| ERR044841-027 | 027 | 2005 | Netherlands | 1 | 2 | ERR044841 |
| ERR044830-027 | 027 | 2006 | Germany | 1 | 2 | ERR044830 |
| ERR044838-027 | 027 | 2006 | Netherlands | 1 | 2 | ERR044838 |
| SRR1519391-LL-027 | 027 | 2004 | UK | 1 | 2 | SRR1519391 |
| ERR044826-027 | 027 | 2005 | Belgium | 1 | 2 | ERR044826 |
| ERR044845-027 | 027 | 2007 | Norway | 1 | 2 | ERR044845 |
| ERR467589-027 | 027 | ? | ? | 1 | 2 | ERR467589 |
| ERR044829-027 | 027 | 2007 | France | 1 | 2 | ERR044829 |
| ERR044844-027 | 027 | 2007 | Netherlands | 1 | 2 | ERR044844 |
| ERR044839-027 | 027 | 2006 | Netherlands | 1 | 2 | ERR044839 |
| ERR044848-027 | 027 | 2008 | Switzerland | 1 | 2 | ERR044848 |
| ERR044834-027 | 027 | 2007 | Luxembourg | 1 | 2 | ERR044834 |
| ERR044833-027 | 027 | 2008 | Ireland | 1 | 2 | ERR044833 |
| ERR044828-027 | 027 | 2009 | Finland | 1 | 2 | ERR044828 |
| ERR026421-027 | 027 | 2006 | UK | 1 | 2 | ERR026421 |
| SRR1519392-LL-029 | 029 | 1995 | UK | 16 | 1 | SRR1519392 |
| SRR593230-029 | 029 | ? | USA | 16 | 1 | SRR593230 |
| ERR833664-031 | 031 | ? | UK | 29 | 1 | ERR833664 |
| SRR593311-031 | 031 | ? | USA | 26 | 1 | SRR593311 |
| SRR1519393-LL-031 | 031 | 1998 | UK | 29 | 1 | SRR1519393 |
| ERR247078-Sang-033 | 033 | 2006 | Australia | 11 | 5 | ERR247078 |
| ERR247077-Sang-033 | 033 | 2009 | UK | 11 | 5 | ERR247077 |
| ERR247076-Sang-033 | 033 | ? | Australia | 11 | 5 | ERR247076 |
| ERR247075-Sang-033 | 033 | 2006 | US | 11 | 5 | ERR247075 |
| ERR247071-Sang-033 | 033 | 2005 | US | 11 | 5 | ERR247071 |
| ERR247070-Sang-033 | 033 | 2007 | US | 11 | 5 | ERR247070 |
| ERR247107-Sang-033 | 033 | 1982 | Australia | 11 | 5 | ERR247107 |
| ERR247106-Sang-033 | 033 | 1980 | Australia | 11 | 5 | ERR247106 |
| ERR247105-Sang-033 | 033 | 2006 | Australia | 11 | 5 | ERR247105 |
| ERR247104-Sang-033 | 033 | 2006 | Australia | 11 | 5 | ERR247104 |
| ERR247103-Sang-033 | 033 | 2006 | Australia | 11 | 5 | ERR247103 |
| ERR2898837-033 | 033 | 1982 | Australia | 11 | 5 | ERR2898837 |
| ERR2898836-033 | 033 | 1980 | Australia | 11 | 5 | ERR2898836 |
| ERR2898835-033 | 033 | 2006 | Australia | 11 | 5 | ERR2898835 |
| ERR2898834-033 | 033 | 2006 | Australia | 11 | 5 | ERR2898834 |
| ERR2898833-033 | 033 | 2006 | Australia | 11 | 5 | ERR2898833 |
| ERR247108-Sang-033 | 033 | 2007 | Australia | 11 | 5 | ERR247108 |
| ERR2898838-033 | 033 | 2007 | Australia | 11 | 5 | ERR2898838 |
| ERR2898928-033 | 033 | 2012 | Australia | 11 | 5 | ERR2898928 |
| ERR2898844-033 | 033 | 2012 | France | 11 | 5 | ERR2898844 |
| ERR247090-Sang-033 | 033 | 2012 | Australia | 11 | 5 | ERR247090 |
| ERR2898920-033 | 033 | 2012 | Australia | 11 | 5 | ERR2898920 |
| ERR247088-Sang-033 | 033 | 2012 | Australia | 11 | 5 | ERR247088 |
| ERR2898923-033 | 033 | 2012 | Australia | 11 | 5 | ERR2898923 |
| ERR2898922-033 | 033 | 2012 | Australia | 11 | 5 | ERR2898922 |
| ERR2898919-033 | 033 | 2012 | Australia | 11 | 5 | ERR2898919 |
| ERR2898885-033 | 033 | 2015 | Australia | 11 | 5 | ERR2898885 |
| ERR2898884-033 | 033 | 2015 | Australia | 11 | 5 | ERR2898884 |
| ERR2898883-033 | 033 | 2015 | Australia | 11 | 5 | ERR2898883 |
| ERR2898882-033 | 033 | 2015 | Australia | 11 | 5 | ERR2898882 |
| ERR2898886-033 | 033 | 2015 | Australia | 11 | 5 | ERR2898886 |
| ERR2898846-033 | 033 | 2012 | France | 11 | 5 | ERR2898846 |
| ERR2898845-033 | 033 | 2013 | France | 11 | 5 | ERR2898845 |
| ERR2898841-033 | 033 | 2012 | France | 11 | 5 | ERR2898841 |
| ERR2898840-033 | 033 | 2013 | Australia | 11 | 5 | ERR2898840 |
| ERR2898831-033 | 033 | 2012 | Australia | 11 | 5 | ERR2898831 |
| ERR2215990-033 | 033 | ? | ? | 11 | 5 | ERR2215990 |
| ERR2898843-033 | 033 | 2011 | France | 11 | 5 | ERR2898843 |
| ERR2898842-033 | 033 | 2012 | France | 11 | 5 | ERR2898842 |
| SRR1519394-LL-033 | 033 | ? | ? | 11 | 5 | SRR1519394 |
| ERR247102-Sang-033 | 033 | ? | Australia | 11 | 5 | ERR247102 |
| ERR171341-Sang-033 | 033 | 2011 | UK | 11 | 5 | ERR171341 |
| ERR247116-Sang-033 | 033 | 2007 | Australia | 11 | 5 | ERR247116 |
| ERR1854839-033 | 033 | 2007 | Australia | 11 | 5 | ERR1854839 |
| ERR2898839-033 | 033 | 2007 | Australia | 11 | 5 | ERR2898839 |
| ERR2898832-033 | 033 | 2012 | Australia | 11 | 5 | ERR2898832 |
| SRR1519395-LL-035 | 035 | ? | ? | 40 | 1 | SRR1519395 |
| 103634-001-033-036 | 036 | ? | ? | 1 | 2 | Not submitted |
| 103634-001-034-036 | 036 | ? | ? | 1 | 2 | Not submitted |
| HNFKWDSXX-103786-001-042-036 | 036 | 2019 | Greece | 1 | 2 | ERR3981060 |
| HNFKWDSXX-103786-001-043-036 | 036 | 2019 | Greece | 1 | 2 | ERR3981061 |
| 103634-001-038-036/198 | 036/198 | ? | Hungary | 1 | 2 | Not submitted |
| 103634-001-041-036/198 | 036/198 | ? | Hungary | 1 | 2 | Not submitted |
| 103634-001-044-036/198 | 036/198 | ? | Hungary | 1 | 2 | Not submitted |
| 103634-001-043-036/198 | 036/198 | ? | Hungary | 1 | 2 | Not submitted |
| 103634-001-037-036/198 | 036/198 | ? | Hungary | 1 | 2 | Not submitted |
| 103634-001-039-036/198 | 036/198 | ? | Hungary | 1 | 2 | Not submitted |
| 103634-001-040-036/198 | 036/198 | ? | Hungary | 1 | 2 | Not submitted |
| 103634-001-042-036/198 | 036/198 | ? | Hungary | 1 | 2 | Not submitted |
| SRR1519396-LL-037 | 037 | 1994 | UK | 98 | 1 | SRR1519396 |
| SRR7852193-039 | 039 | ? | UK | ? (unknown) | ? | SRR7852193 |
| SRR7852194-039 | 039 | ? | UK | ? (unknown) | ? | SRR7852194 |
| SRR1519397-LL-039 | 039 | ? | ? | 26 | 1 | SRR1519397 |
| SRR1519398-LL-040 | 040 | ? | ? | 121 | 4 | SRR1519398 |
| SRR1519399-LL-042 | 042 | 1998 | UK | 6 | 1 | SRR1519399 |
| SRR1519400-LL-043 | 043 | 1980 | New Zealand | 103 | 1 | SRR1519400 |
| ERR247084-Sang-045 | 045 | 2009 | Australia | 11 | 5 | ERR247084 |
| ERR256923-Sang-045 | 045 | 2010 | UK | 11 | 5 | ERR256923 |
| ERR256920-Sang-045 | 045 | 2008 | UK | 11 | 5 | ERR256920 |
| ERR247091-Sang-045 | 045 | 2005 | Australia | 11 | 5 | ERR247091 |
| ERR247114-Sang-045 | 045 | 2006 | Australia | 11 | 5 | ERR247114 |
| ERR247117-Sang-045 | 045 | 2007 | Australia | 11 | 5 | ERR247117 |
| ERR257024-Sang-045 | 045 | 2012 | Sweden | 161 | 5 | ERR257024 |
| ERR171222-Sang-045 | 045 | 2009 | UK | 11 | 5 | ERR171222 |
| ERR171342-Sang-045 | 045 | 2011 | UK | 11 | 5 | ERR171342 |
| SRR1519401-LL-045 | 045 | ? | ? | 11 | 5 | SRR1519401 |
| ERR126280-045 | 045 | ? | ? | 11 | 5 | ERR126280 |
| ERR171355-Sang-045 | 045 | 2011 | Netherlands | 161 | 5 | ERR171355 |
| ERR171354-Sang-045 | 045 | 2011 | Netherlands | 161 | 5 | ERR171354 |
| ERR256921-Sang-045 | 045 | 2009 | UK | 11 | 5 | ERR256921 |
| ERR256924-Sang-045 | 045 | 2010 | UK | 11 | 5 | ERR256924 |
| SRR593309-046 | 046 | ? | USA | 35 | 1 | SRR593309 |
| SRR1519402-LL-046 | 046 | ? | ? | 35 | 1 | SRR1519402 |
| ERR171363-Sang-047 | 047 | 2009 | UK | 37 | 4 | ERR171363 |
| SRR1519403-LL-047 | 047 | 2007 | UK | 37 | 4 | SRR1519403 |
| SRR1519404-050 | 050 | 2007 | UK | 18 | 1 | SRR1519404 |
| SRR1519405-LL-051 | 051 | 1996 | UK | 101 | 1 | SRR1519405 |
| SRR1519406-LL-053 | 053 | 2006 | USA | 63 | 1 | SRR1519406 |
| SRR1519407-LL-054 | 054 | ? | ? | 43 | 1 | SRR1519407 |
| SRR593228-054 | 054 | ? | USA | 43 | 1 | SRR593228 |
| SRR1519408-LL-055 | 055 | 1998 | UK | 3 | 1 | SRR1519408 |
| SRR593371-056 | 056 | ? | USA | 58 | 1 | SRR593371 |
| SRR593385-056 | 056 | ? | USA | 34 | 1 | SRR593385 |
| SRR1519409-LL-056 | 056 | ? | ? | 34 | 1 | SRR1519409 |
| SRR1519410-LL-057 | 057 | 2008 | UK | 55 | 1 | SRR1519410 |
| ERR171364-Sang-058 | 058 | 2009 | UK | 96 | 3 | ERR171364 |
| SRR1519411-LL-058 | 058 | 1997 | UK | 96 | 3 | SRR1519411 |
| ERR126275-058 | 058 | 2008 | UK | 96 | 3 | ERR126275 |
| ERR125966-Sang-060 | 060 | 2009 | UK | 87 | 4 | ERR125966 |
| SRR1519412-LL-060 | 060 | ? | ? | 38 | 4 | SRR1519412 |
| SRR1519413-LL-062 | 062 | 2008 | UK | 44 | 1 | SRR1519413 |
| SRR593211-062 | 062 | ? | USA | 44 | 1 | SRR593211 |
| SRR1519414-LL-063 | 063 | ? | ? | 5 | 3 | SRR1519414 |
| SRR1519415-LL-064 | 064 | ? | ? | 33 | 1 | SRR1519415 |
| ERR171335-066 | 066 | ? | UK | 11 | 5 | ERR171335 |
| ERR171343-Sang-066 | 066 | 2010 | UK | 11 | 5 | ERR171343 |
| SRR1519416-LL-066 | 066 | ? | UK | 11 | 5 | SRR1519416 |
| SRR1519417-LL-067 | 067 | ? | ? | 27 | 1 | SRR1519417 |
| SRR1519418-LL-068 | 068 | ? | ? | 127 | 4 | SRR1519418 |
| ERR126286-069 | 069 | 2009 | UK | 5 | 3 | ERR126286 |
| SRR7852179-070 | 070 | ? | UK | 99 | 1 | SRR7852179 |
| SRR1519419-LL-070 | 070 | ? | ? | 55 | 1 | SRR1519419 |
| SRR1519420-LL-072 | 072 | ? | ? | 3 | 1 | SRR1519420 |
| SRR1519421-LL-075 | 075 | ? | ? | 95 | 2 | SRR1519421 |
| SRR1519422-LL-076 | 076 | 1997 | UK | 2 | 1 | SRR1519422 |
| ERR3278159-076 | 076 | ? | Spain | 2 | 1 | ERR3278159 |
| SRR6042350-077 | 077 | 2010 | USA | 3 | 1 | SRR6042350 |
| ERR2898875-078 | 078 | 2012 | Australia | 11 | 5 | ERR2898875 |
| ERR171325-078 | 078 | 2008 | Netherlands | 11 | 5 | ERR171325 |
| ERR171328-078 | 078 | 2009 | Netherlands | 11 | 5 | ERR171328 |
| ERR171329-078 | 078 | 2010 | Netherlands | 11 | 5 | ERR171329 |
| ERR257065-078-Pig-P12 | 078 | 2011 | Netherlands | 11 | 5 | ERR257065 |
| ERR257061-078-Farm-P4 | 078 | 2011 | Netherlands | 11 | 5 | ERR257061 |
| ERR257046-078-Pig-P4 | 078 | 2011 | Netherlands | 11 | 5 | ERR257046 |
| ERR257045-078-Pig-P9 | 078 | 2011 | Netherlands | 11 | 5 | ERR257045 |
| ERR257044-078-Farm-P12 | 078 | 2011 | Netherlands | 11 | 5 | ERR257044 |
| ERR257050-078-Farm-P8 | 078 | 2011 | Netherlands | 11 | 5 | ERR257050 |
| SRR7308938-078 | 078 | 2013 | Germany | 11 | 5 | SRR7308938 |
| ERR171322-078 | 078 | 2008 | Netherlands | 11 | 5 | ERR171322 |
| ERR171332-078 | 078 | 2011 | Netherlands | 11 | 5 | ERR171332 |
| ERR171337-078 | 078 | 2002 | Netherlands | 11 | 5 | ERR171337 |
| ERR171338-078 | 078 | 2002 | Netherlands | 11 | 5 | ERR171338 |
| ERR257070-078-Pig-P11 | 078 | 2011 | Netherlands | 11 | 5 | ERR257070 |
| ERR257048-078-Pig-P10 | 078 | 2011 | Netherlands | 11 | 5 | ERR257048 |
| ERR257051-078-Pig-P6 | 078 | 2011 | Netherlands | 11 | 5 | ERR257051 |
| CD05_A_078 | 078 | 2018 | Netherlands | 11 | 5 | ERS6671669 (This study) |
| CD07_A_078 | 078 | 2018 | Netherlands | 11 | 5 | ERS6671670 (This study) |
| CD08_A_078 | 078 | 2018 | Netherlands | 11 | 5 | ERS6671671 (This study) |
| ERR171317-078 | 078 | 2007 | Netherlands | 11 | 5 | ERR171317 |
| ERR257063-078-Pig-P2 | 078 | 2011 | Netherlands | 11 | 5 | ERR257063 |
| ERR257049-078-Farm-P2 | 078 | 2011 | Netherlands | 11 | 5 | ERR257049 |
| ERR257053-078-Pig-P3 | 078 | 2011 | Netherlands | 11 | 5 | ERR257053 |
| CD3-2_DMS27440-V-078 | 078 | ? | China | 11 | 5 | Not submitted |
| ERR257058-078-Pig-UP | 078 | 2011 | Netherlands | 11 | 5 | ERR257058 |
| SRR7308955-078 | 078 | 2013 | Greece | 11 | 5 | SRR7308955 |
| SRR7308972-078 | 078 | 2012 | France | 11 | 5 | SRR7308972 |
| SRR7309155-078 | 078 | 2013 | Italy | 11 | 5 | SRR7309155 |
| CD09_A_078 | 078 | 2018 | Netherlands | 11 | 5 | ERS6671672 (This study) |
| CD10_A_078 | 078 | 2018 | Netherlands | 11 | 5 | ERS6671673 (This study) |
| SRR7309171-078 | 078 | 2013 | Portugal | 11 | 5 | SRR7309171 |
| SRR7309098-078 | 078 | 2013 | Italy | 11 | 5 | SRR7309098 |
| SRR7308946-078 | 078 | 2013 | Germany | 11 | 5 | SRR7308946 |
| SRR7308846-078 | 078 | 2013 | Germany | 11 | 5 | SRR7308846 |
| SRR1519423-LL-078 | 078 | ? | ? | 11 | 5 | SRR1519423 |
| ERR2898853-078 | 078 | 2016 | ERR2898853 | 11 | 5 | ERR2898853 |
| ERR2898848-078 | 078 | 2010 | Australia | 11 | 5 | ERR2898848 |
| ERR027342-078 | 078 | 2007 | Ireland | 11 | 5 | ERR027342 |
| SRR7308986-078 | 078 | 2013 | France | 11 | 5 | SRR7308986 |
| SRR7308758-078 | 078 | 2013 | Ireland | 11 | 5 | SRR7308758 |
| SRR7309059-078 | 078 | 2013 | UK | 11 | 5 | SRR7309059 |
| SRR7309228-078 | 078 | 2013 | Italy | 11 | 5 | SRR7309228 |
| ERR171331-078 | 078 | 2010 | Netherlands | 11 | 5 | ERR171331 |
| ERR171309-078 | 078 | 2006 | Netherlands | 11 | 5 | ERR171309 |
| ERR257068-078-Pig-UP | 078 | 2011 | Netherlands | 11 | 5 | ERR257068 |
| ERR171313-078 | 078 | 2007 | Netherlands | 11 | 5 | ERR171313 |
| ERR171352-078-Pig-UP | 078 | 2009 | Netherlands | 11 | 5 | ERR171352 |
| ERR171353-078-Pig-UP | 078 | 2009 | Netherlands | 11 | 5 | ERR171353 |
| ERR171316-078 | 078 | 2007 | Netherlands | 11 | 5 | ERR171316 |
| ERR171319-078 | 078 | 2007 | Netherlands | 11 | 5 | ERR171319 |
| ERR171320-078 | 078 | 2008 | Netherlands | 11 | 5 | ERR171320 |
| ERR257059-078-Farm-UP | 078 | 2011 | Netherlands | 11 | 5 | ERR257059 |
| ERR171323-078 | 078 | 2008 | Netherlands | 11 | 5 | ERR171323 |
| ERR171326-078 | 078 | 2009 | Netherlands | 11 | 5 | ERR171326 |
| ERR171330-078 | 078 | 2010 | Netherlands | 11 | 5 | ERR171330 |
| ERR171339-078 | 078 | 2002 | Netherlands | 11 | 5 | ERR171339 |
| ERR171343-078 | 078 | 2010 | Netherlands | 11 | 5 | ERR171343 |
| ERR171346-078 | 078 | 2007 | Netherlands | ? | ? | ERR171346 |
| ERR171349-078 | 078 | 2009 | Netherlands | 11 | 5 | ERR171349 |
| ERR257066-078-Farm-P7 | 078 | 2011 | Netherlands | 11 | 5 | ERR257066 |
| ERR257064-078-Pig-P7 | 078 | 2011 | Netherlands | 11 | 5 | ERR257064 |
| ERR171310-078 | 078 | 2006 | Netherlands | 11 | 5 | ERR171310 |
| ERR171312-078 | 078 | 2007 | Netherlands | 11 | 5 | ERR171312 |
| ERR171314-078 | 078 | 2007 | Netherlands | 11 | 5 | ERR171314 |
| ERR171321-078 | 078 | 2008 | Netherlands | 11 | 5 | ERR171321 |
| ERR171327-078 | 078 | 2009 | Netherlands | 11 | 5 | ERR171327 |
| ERR403717-078 | 078 | 2008 | Netherlands | 11 | 5 | ERR403717 |
| ERR257072-078-Farm-P5 | 078 | 2011 | Netherlands | 11 | 5 | ERR257072 |
| ERR257071-78-Pig-P5 | 078 | 2011 | Netherlands | 11 | 5 | ERR257071 |
| ERR257069-078-Farm-P6 | 078 | 2011 | Netherlands | 11 | 5 | ERR257069 |
| ERR257056-078-Farm-P10 | 078 | 2011 | Netherlands | 11 | 5 | ERR257056 |
| ERR257055-078-Pig-P8 | 078 | 2011 | Netherlands | 11 | 5 | ERR257055 |
| ERR257054-078-Farm-UP | 078 | 2011 | Netherlands | 11 | 5 | ERR257054 |
| ERR171356-078-Farm-P1 | 078 | 2011 | Netherlands | 11 | 5 | ERR171356 |
| ERR257047-078-Farm-P11 | 078 | 2011 | Netherlands | 11 | 5 | ERR257047 |
| ERR257060-078-Farm-P9 | 078 | 2011 | Netherlands | 11 | 5 | ERR257060 |
| ERR257062-078-Farm-UP | 078 | 2011 | Netherlands | 11 | 5 | ERR257062 |
| SRR7308726-078 | 078 | 2013 | UK | 11 | 5 | SRR7308726 |
| SRR7308912-078 | 078 | 2013 | Ireland | 11 | 5 | SRR7308912 |
| SRR7308920-078 | 078 | 2013 | Spain | 11 | 5 | SRR7308920 |
| SRR7309151-078 | 078 | 2013 | Germany | 11 | 5 | SRR7309151 |
| CD11_A_078 | 078 | 2018 | Netherlands | 11 | 5 | ERS6671674 (This study) |
| SRR7308895-078 | 078 | 2013 | Germany | 11 | 5 | SRR7308895 |
| ERR171311-078 | 078 | 2007 | Netherlands | 11 | 5 | ERR171311 |
| ERR171315-078 | 078 | 2007 | Netherlands | 11 | 5 | ERR171315 |
| ERR171318-078 | 078 | 2007 | Netherlands | 11 | 5 | ERR171318 |
| ERR171334-078 | 078 | 2009 | Netherlands | 11 | 5 | ERR171334 |
| ERR171336-078 | 078 | 2002 | Netherlands | 11 | 5 | ERR171336 |
| ERR171347-078-Pig-UP | 078 | 2002 | Netherlands | 11 | 5 | ERR171347 |
| ERR171348-078-Pig-UP | 078 | 2009 | Netherlands | 11 | 5 | ERR171348 |
| SRR7308809-078 | 078 | 2013 | Germany | 11 | 5 | SRR7308809 |
| SRR7308838-078 | 078 | 2013 | Portugal | 11 | 5 | SRR7308838 |
| ERR171357-078-Pig-P1 | 078 | 2011 | Netherlands | 11 | 5 | ERR171357 |
| SRR7308822-078 | 078 | 2013 | Italy | 11 | 5 | SRR7308822 |
| SRR7309025-078 | 078 | 2013 | Italy | 11 | 5 | SRR7309025 |
| SRR7309049-078 | 078 | 2013 | Portugal | 11 | 5 | SRR7309049 |
| SRR1519424-LL-079 | 079 | 1994 | ? | 126 | 1 | SRR1519424 |
| SRR1519425-LL-081 | 081 | 1996 | UK | 9 | 1 | SRR1519425 |
| SRR7852189-081 | 081 | ? | UK | 9 | 1 | SRR7852189 |
| SRR7852192-081 | 081 | ? | UK | 9 | 1 | SRR7852192 |
| SRR1519426-LL-083 | 083 | 1997 | UK | 59 | 1 | SRR1519426 |
| SRR1519427-LL-084 | 084 | 1995 | UK | 48 | 1 | SRR1519427 |
| SRR593202-084 | 084 | ? | USA | 17 | 1 | SRR593202 |
| ERR125915-Sang-085 | 085 | 2009 | UK | 39 | 4 | ERR125915 |
| SRR1519428-LL-085 | 085 | ? | ? | 39 | 4 | SRR1519428 |
| ERR2216001-087 | 087 | ? | ? | 46 | 1 | ERR2216001 |
| SRR1519429-LL-087 | 087 | ? | ? | 46 | 1 | SRR1519429 |
| SRR1519430-LL-088 | 088 | 1997 | UK | 39 | 4 | SRR1519430 |
| SRR1519431-LL-095 | 095 | 1995 | ? | 2 | 1 | SRR1519431 |
| ERR2215983-102 | 102 | ? | ? | 24 | 1 | ERR2215983 |
| SRR593300-103 | 103 | ? | USA | 53 | 1 | SRR593300 |
| ERR2215975-103 | 103 | ? | ? | 53 | 1 | ERR2215975 |
| ERR3276437-106 | 106 | ? | ERR3276437 | 42 | 1 | ERR3276437 |
| ERR3276441-106 | 106 | ? | Spain | 42 | 1 | ERR3276441 |
| ERR3276506-106 | 106 | ? | Spain | 42 | 1 | ERR3276506 |
| ERR3278163-106 | 106 | ? | Spain | 42 | 1 | ERR3278163 |
| ERR3278167-106 | 106 | ? | Spain | 42 | 1 | ERR3278167 |
| ERR3288184-106 | 106 | ? | Spain | 42 | 1 | ERR3288184 |
| ERR3288190-106 | 106 | ? | Spain | 42 | 1 | ERR3288190 |
| ERR3288325-106 | 106 | ? | Spain | 42 | 1 | ERR3288325 |
| ERR3288336-106 | 106 | ? | Spain | 42 | 1 | ERR3288336 |
| ERR3288338-106 | 106 | ? | Spain | 42 | 1 | ERR3288338 |
| ERR3289201-106 | 106 | ? | Spain | 42 | 1 | ERR3289201 |
| ERR3289202-106 | 106 | ? | Spain | 42 | 1 | ERR3289202 |
| ERR3289205-106 | 106 | ? | Spain | 42 | 1 | ERR3289205 |
| ERR3289207-106 | 106 | ? | Spain | 42 | 1 | ERR3289207 |
| ERR3289209-106 | 106 | ? | Spain | 42 | 1 | ERR3289209 |
| ERR3289212-106 | 106 | ? | Spain | 42 | 1 | ERR3289212 |
| ERR3299518-106 | 106 | ? | Spain | 42 | 1 | ERR3299518 |
| ERR3288471-106 | 106 | ? | Spain | 42 | 1 | ERR3288471 |
| ERR3289206-106 | 106 | ? | Spain | 42 | 1 | ERR3289206 |
| ERR3276440-106 | 106 | ? | Spain | 42 | 1 | ERR3276440 |
| ERR3299515-106 | 106 | ? | Spain | 42 | 1 | ERR3299515 |
| ERR3288467-106 | 106 | ? | Spain | 42 | 1 | ERR3288467 |
| ERR3288205-106 | 106 | ? | Spain | 42 | 1 | ERR3288205 |
| ERR3288577-106 | 106 | ? | Spain | 42 | 1 | ERR3288577 |
| ERR3288578-106 | 106 | ? | Spain | 42 | 1 | ERR3288578 |
| ERR3289203-106 | 106 | ? | Spain | 42 | 1 | ERR3289203 |
| ERR3296455-106 | 106 | ? | Spain | 42 | 1 | ERR3296455 |
| ERR3299510-106 | 106 | ? | Spain | 42 | 1 | ERR3299510 |
| SRR1519432-LL-106 | 106 | 1997 | UK | 42 | 1 | SRR1519432 |
| SRR6042360-106/174 | 106/174 | 2010 | USA | 36 | 1 | SRR6042360 |
| SRR6042371-106/174 | 106/174 | 2010 | USA | 42 | 1 | SRR6042371 |
| SRR593379-111 | 111 | ? | USA | 123 | 2 | SRR593379 |
| SRR1519433-LL-115 | 115 | 1997 | UK | 3 | 1 | SRR1519433 |
| SRR1519434-LL-117 | 117 | 1996 | Poland | 54 | 1 | SRR1519434 |
| SRR1519435-LL-118 | 118 | 2007 | UK | 42 | 1 | SRR1519435 |
| SRR1519436-LL-122 | 122 | ? | ? | 116 | 2 | SRR1519436 |
| ERR247101-Sang-126 | 126 | ? | Australia | 258 | 5 | ERR247101 |
| ERR256918-Sang-126 | 126 | 2011 | UK | 11 | 5 | ERR256918 |
| ERR247083-Sang-126 | 126 | 2009 | Australia | 258 | 5 | ERR247083 |
| ERR2898771-126 | 126 | 2002 | Italy | 11 | 5 | ERR2898771 |
| ERR2898789-126 | 126 | 2012 | Taiwan | 11 | 5 | ERR2898789 |
| ERR2898749-126 | 126 | 2013 | Australia | 11 | 5 | ERR2898749 |
| SRR1519437-LL-126 | 126 | ? | ? | 11 | 5 | SRR1519437 |
| ERR256915-Sang-126 | 126 | 2010 | UK | 11 | 5 | ERR256915 |
| ERR257010-Sang-126 | 126 | 2010 | Belgium | 11 | 5 | ERR257010 |
| ERR257012-Sang-126 | 126 | 2010 | Belgium | 11 | 5 | ERR257012 |
| ERR257015-Sang-126 | 126 | 2010 | Belgium | 11 | 5 | ERR257015 |
| ERR257016-Sang-126 | 126 | 2010 | Belgium | 11 | 5 | ERR257016 |
| ERR257018-Sang-126 | 126 | 2010 | Belgium | 11 | 5 | ERR257018 |
| ERR257019-Sang-126 | 126 | 2010 | Belgium | 11 | 5 | ERR257019 |
| ERR257021-Sang-126 | 126 | 2010 | Belgium | 11 | 5 | ERR257021 |
| ERR257022-Sang-126 | 126 | 2010 | Belgium | 11 | 5 | ERR257022 |
| ERR2898787-126 | 126 | 2011 | Taiwan | 11 | 5 | ERR2898787 |
| ERR2898786-126 | 126 | ? | Spain | 11 | 5 | ERR2898786 |
| ERR2898746-126 | 126 | 2011 | Australia | 11 | 5 | ERR2898746 |
| ERR2898745-126 | 126 | 2011 | Australia | 11 | 5 | ERR2898745 |
| ERR2898783-126 | 126 | 2010 | Italy | 11 | 5 | ERR2898783 |
| ERR2898748-126 | 126 | 2012 | Australia | 11 | 5 | ERR2898748 |
| ERR2898747-126 | 126 | 2012 | Australia | 11 | 5 | ERR2898747 |
| ERR2898769-126 | 126 | ? | Canada | 11 | 5 | ERR2898769 |
| ERR2898750-126 | 126 | 2012 | Australia | 11 | 5 | ERR2898750 |
| ERR2898772-126 | 126 | 2013 | Italy | 11 | 5 | ERR2898772 |
| ERR2898791-126 | 126 | ? | USA | 11 | 5 | ERR2898791 |
| ERR2898782-126 | 126 | 2008 | Italy | ? | ? | ERR2898782 |
| ERR256916-Sang-126 | 126 | 2010 | UK | 11 | 5 | ERR256916 |
| ERR2898818-127 | 127 | 2008 | Australia | 11 | 5 | ERR2898818 |
| ERR2898900-127 | 127 | 2012 | Australia | 11 | 5 | ERR2898900 |
| ERR2898860-127 | 127 | 2007 | Australia | 11 | 5 | ERR2898860 |
| ERR2898807-127 | 127 | 2013 | Australia | 11 | 5 | ERR2898807 |
| ERR2898911-127 | 127 | 2013 | Australia | 11 | 5 | ERR2898911 |
| ERR2898801-127 | 127 | 2012 | Australia | 11 | 5 | ERR2898801 |
| ERR2898802-127 | 127 | 2014 | Australia | 11 | 5 | ERR2898802 |
| ERR247118-Sang-127 | 127 | 2005 | Australia | 11 | 5 | ERR247118 |
| ERR2898820-127 | 127 | 2005 | Australia | 11 | 5 | ERR2898820 |
| ERR247093-Sang-127 | 127 | 2005 | Australia | 11 | 5 | ERR247093 |
| ERR2898799-127 | 127 | 2011 | Australia | 11 | 5 | ERR2898799 |
| ERR2898823-127 | 127 | 2010 | Japan | 11 | 5 | ERR2898823 |
| ERR2898825-127 | 127 | 2010 | Japan | 11 | 5 | ERR2898825 |
| ERR2898824-127 | 127 | 2010 | Japan | 11 | 5 | ERR2898824 |
| ERR2898828-127 | 127 | 2011 | Taiwan | 11 | 5 | ERR2898828 |
| ERR2898829-127 | 127 | 2012 | Taiwan | 11 | 5 | ERR2898829 |
| ERR2898898-127 | 127 | 2009 | Australia | 11 | 5 | ERR2898898 |
| ERR171360-Sang-130 | 130 | 2010 | UK | 158 | 4 | ERR171360 |
| SRR1519438-131 | 131 | 2001 | Kuwait | 122 | ? | SRR1519438 |
| SRR593375-137 | 137 | ? | USA | 4 | 1 | SRR593375 |
| SRR1519439-LL-153 | 153 | 2003 | Netherlands | 32 | 2 | SRR1519439 |
| SRR1519440-169 | 169 | 2005 | UK | 13 | 1 | SRR1519440 |
| ERR171361-Sang-172 | 172 | 2010 | UK | 159 | 4 | ERR171361 |
| SRR1519441-LL-174 | 174 | 2007 | UK | 42 | 1 | SRR1519441 |
| SRR7852188-174 | 174 | ? | UK | 6 | 1 | SRR7852188 |
| SRR7308767-176 | 176 | 2013 | Germany | 1 | 2 | SRR7308767 |
| SRR7309085-176 | 176 | 2013 | Germany | 1 | 2 | SRR7309085 |
| SRR7308880-176 | 176 | 2013 | Germany | 1 | 2 | SRR7308880 |
| SRR7308660-176 | 176 | 2013 | Czech Republic | 1 | 2 | SRR7308660 |
| SRR7308694-176 | 176 | 2013 | Czech Republic | 1 | 2 | SRR7308694 |
| SRR7309134-176 | 176 | 2013 | Czech Republic | 1 | 2 | SRR7309134 |
| SRR7309148-176 | 176 | 2013 | Czech Republic | 1 | 2 | SRR7309148 |
| SRR7309197-176 | 176 | 2013 | Czech Republic | 1 | 2 | SRR7309197 |
| SRR7308825-176 | 176 | 2013 | Czech Republic | 1 | 2 | SRR7308825 |
| SRR7308625-176 | 176 | 2013 | Czech Republic | 1 | 2 | SRR7308625 |
| SRR7309097-176 | 176 | 2013 | Czech Republic | 1 | 2 | SRR7309097 |
| SRR7308956-176 | 176 | 2013 | Czech Republic | 1 | 2 | SRR7308956 |
| SRR7308940-176 | 176 | 2013 | Czech republic | 1 | 2 | SRR7308940 |
| SRR7308771-176 | 176 | 2013 | Czech republic | 1 | 2 | SRR7308771 |
| SRR7308985-176 | 176 | 2013 | Germany | 1 | 2 | SRR7308985 |
| SRR7308631-176 | 176 | 2013 | Germany | 1 | 2 | SRR7308631 |
| HNFKWDSXX-103786-001-033-181 | 181 | 2019 | Greece | 1 | 2 | ERR3981062 |
| HNFKWDSXX-103786-001-034-181 | 181 | 2019 | Greece | 1 | 2 | ERR3981063 |
| HNFKWDSXX-103786-001-035-181 | 181 | 2019 | Greece | 1 | 2 | ERR3981064 |
| HNFKWDSXX-103786-001-036-181 | 181 | 2019 | Greece | 1 | 2 | ERR3981065 |
| HNFKWDSXX-103786-001-037-181 | 181 | 2019 | Greece | 1 | 2 | ERR3981066 |
| HNFKWDSXX-103786-001-038-181 | 181 | 2019 | Greece | 1 | 2 | ERR3981067 |
| HNFKWDSXX-103786-001-039-181 | 181 | 2019 | Greece | 1 | 2 | ERR3981068 |
| HNFKWDSXX-103786-001-040-181 | 181 | 2019 | Greece | 1 | 2 | ERR3981069 |
| HNFKWDSXX-103786-001-041-181 | 181 | 2019 | Greece | 1 | 2 | ERR3981070 |
| HNFKWDSXX-103786-001-044-181 | 181 | 2019 | Greece | 1 | 2 | ERR3981071 |
| HNFKWDSXX-103786-001-045-181 | 181 | 2019 | Greece | 1 | 2 | ERR3981072 |
| HNFKWDSXX-103786-001-046-181 | 181 | 2019 | Greece | 1 | 2 | ERR3981073 |
| HNFKWDSXX-103786-001-047-181 | 181 | 2019 | Greece | 1 | 2 | ERR3981074 |
| HNFKWDSXX-103786-001-048-181 | 181 | 2019 | Greece | 1 | 2 | ERR3981075 |
| HNFKWDSXX-103786-001-049-181 | 181 | 2019 | Greece | 1 | 2 | ERR3981076 |
| Isolate1-Leeds | 181 | ? | Romania | 1 | 2 | ERR3981079 |
| Isolate2-Leeds | 181 | ? | Romania | 1 | 2 | ERR3981080 |
| SRR1519442-LL-198 | 198 | 2006 | UK | 1 | 2 | SRR1519442 |
| 103634-001-035-198 | 198 | ? | ? | 1 | 2 | Not submitted |
| ERR833670-220 | 220 | ? | UK | 2 | 1 | ERR833670 |
| ERR833669-220 | 220 | ? | UK | 2 | 1 | ERR833669 |
| SRR7852190-225 | 225 | ? | UK | 58 | 1 | SRR7852190 |
| ERR247085-Sang-237 | 237 | 2009 | Australia | 167 | 5 | ERR247085 |
| ERR247082-Sang-237 | 237 | 2008 | Australia | 167 | 5 | ERR247082 |
| ERR1854833-238 | 238 | 2007 | Australia | 169 | 5 | ERR1854833 |
| ERR1854841-239 | 239 | 2005 | Australia | 168 | 5 | ERR1854841 |
| SRR2751309-244 | 244 | ? | Australia | 41 | 2 | SRR2751309 |
| SRR2751311-244 | 244 | ? | Australia | 41 | 2 | SRR2751311 |
| SRR2751302-Sang-244 | 244 | ? | Australia | 41 | 2 | SRR2751302 |
| SRR2751305-244 | 244 | ? | Australia | 41 | 2 | SRR2751305 |
| SRR2751307-244 | 244 | ? | Australia | 41 | 2 | SRR2751307 |
| SRR2751308-244 | 244 | ? | Australia | 41 | 2 | SRR2751308 |
| SRR2751310-244 | 244 | ? | Australia | 41 | 2 | SRR2751310 |
| SRR2751293-244 | 244 | ? | Australia | 41 | 2 | SRR2751293 |
| ERR2215996-Clade2-244 | 244 | ? | ? | 41 | 2 | ERR2215996 |
| ERR2215977-Clade2-251 | 251 | ? | ? | 231 | 2 | ERR2215977 |
| ERR1347091-251 | 251 | ? | Australia | 231 | 2 | ERR1347091 |
| ERR1347090-251 | 251 | ? | Australia | 231 | 2 | ERR1347090 |
| ERR1347089-251 | 251 | ? | Australia | 231 | 2 | ERR1347089 |
| ERR1347088-251 | 251 | ? | Australia | 231 | 2 | ERR1347088 |
| ERR2898847-288 | 288 | 2006 | Australia | 11 | 5 | ERR2898847 |
| ERR2898924-288 | 288 | 2013 | Australia | 11 | 5 | ERR2898924 |
| SRR7308916-356 | 356 | 2013 | Italy | 17 | 1 | SRR7308916 |
| SRR7309178-356 | 356 | 2013 | Italy | 17 | 1 | SRR7309178 |
| SRR7309102-356 | 356 | 2013 | Italy | ? | ? | SRR7309102 |
| SRR7309088-356 | 356 | 2013 | Italy | 17 | 1 | SRR7309088 |
| SRR7309077-356 | 356 | 2013 | Italy | 17 | 1 | SRR7309077 |
| SRR7309044-356 | 356 | 2013 | Italy | 17 | 1 | SRR7309044 |
| SRR7309041-356 | 356 | 2013 | Italy | 17 | 1 | SRR7309041 |
| SRR7309022-356 | 356 | 2013 | Italy | 17 | 1 | SRR7309022 |
| SRR7308855-356 | 356 | 2013 | Italy | 17 | 1 | SRR7308855 |
| SRR7308803-356 | 356 | 2013 | Italy | 17 | 1 | SRR7308803 |
| SRR7308786-356 | 356 | 2013 | Italy | 17 | 1 | SRR7308786 |
| SRR7308719-356 | 356 | 2013 | Italy | 17 | 1 | SRR7308719 |
| SRR7308708-356 | 356 | 2013 | Italy | 17 | 1 | SRR7308708 |
| ERR3293604-404 | 404 | ? | Spain | 110 | 1 | ERR3293604 |
| ERR3293601-404 | 404 | ? | Spain | 110 | 1 | ERR3293601 |
| ERR3289211-404 | 404 | ? | Spain | 110 | 1 | ERR3289211 |
| ERR3288472-404 | 404 | ? | Spain | 110 | 1 | ERR3288472 |
| ERR3289197-404 | 404 | ? | Spain | 110 | 1 | ERR3289197 |
| ERR3288469-404 | 404 | ? | Spain | 110 | 1 | ERR3288469 |
| ERR3288324-404 | 404 | ? | Spain | 110 | 1 | ERR3288324 |
| ERR3288327-404 | 404 | ? | Spain | 110 | 1 | ERR3288327 |
| ERR3288177-404 | 404 | ? | Spain | 110 | 1 | ERR3288177 |
| ERR3288176-404 | 404 | ? | Spain | 110 | 1 | ERR3288176 |
| ERR3278162-404 | 404 | ? | Spain | 110 | 1 | ERR3278162 |
| ERR3274941-404 | 404 | ? | Spain | 110 | 1 | ERR3274941 |
| ERR3288348-404 | 404 | ? | Spain | 110 | 1 | ERR3288348 |
| ERR1854837-585 | 585 | 1998 | ? | 164 | 5 | ERR1854837 |
| ERR1854838-586 | 586 | 2007 | Australia | 167 | 5 | ERR1854838 |

